# Nucleotide excision repair hotspots and coldspots of UV-induced DNA damage in the human genome

**DOI:** 10.1101/2020.04.16.045369

**Authors:** Yuchao Jiang, Wentao Li, Laura A Lindsey-Boltz, Yuchen Yang, Yun Li, Aziz Sancar

## Abstract

We recently developed high-throughput sequencing approaches, eXcision Repair sequencing (XR-seq) and Damage-seq, to generate genome-wide mapping of DNA excision repair and damage formation, respectively, with single-nucleotide resolution. Here, we used time-course XR-seq data to profile UV-induced excision repair dynamics, paired with Damage-seq data to quantify the overall induced DNA damage. We identified genome-wide repair hotspots exhibiting high-level nucleotide excision repair immediately after UV irradiation. We show that such repair hotspots do not result from hypersensitivity to DNA damage, and are thus not damage hotspots. We find that the earliest repair occurs preferentially in promoters and enhancers from open-chromatin regions. The repair hotspots are also significantly enriched for frequently interacting regions and super-enhancers, both of which are themselves hotspots for local chromatin interactions. Further interrogation of chromatin organization to include DNA replication timing allows us to conclude that early-repair hotspots are enriched for early-replication domains. Collectively, we report genome-wide early-repair hotspots of UV-induced damage, in association with chromatin states and epigenetic compartmentalization of the human genome.

## INTRODUCTION

UV in sunlight is a known mutagen and causative agent of skin cancer that induces DNA lesions such as cyclobutane pyrimidine dimers (CPDs) and pyrimidine-pyrimidone (6-4) photoproducts [(6-4)PPs] (1,2). In humans, both damage types are solely repaired by nucleotide excision repair (excision repair), which removes DNA lesions by dual incisions bracketing modified bases and fills and seals the resulting gap by DNA synthesis and ligation (3). Excision repair consists of two pathways, global repair and transcription-coupled repair (4,5), that differ primarily in the damage recognition step (3-6). For UV-induced DNA damage, CPD repair is highly associated with transcription, specifically with the transcribed strand, while (6-4)PP repair is uniformly distributed throughout the genome (7).

Recently, we developed a high-throughput approach, excision repair sequencing (XR-seq), to isolate the oligonucleotides excised by excision repair and subject them to next-generation sequencing (7). This method has allowed for genome-wide mapping of UV-induced excision repair with single-nucleotide resolution in human (7), Lemur (8), mouse (9), *Drosophila melanogaster* (10), *Saccharomyces cerevisiae* (11), *Escherichia coli* (12), and *Arabidopsis thaliana* (13). We further developed another next-generation sequencing method, Damage-seq (14), to generate genome-wide mapping of UV damage formation with single-nucleotide resolution (15).

Using the combined UV damage maps and repair maps, it has been shown that the induced DNA damage is uniformly distributed throughout the genome and that the overall effect of damage in the genome is primarily driven, not by damage formation, but by repair efficiency (14,15). Indeed, different repair efficiencies across the genome have been previously reported, which are impacted by multiple factors, including transcription, chromatin state and structure, regulatory protein binding to DNA, and posttranscriptional modification of histones (7,15-22). Existing studies, however, have been focused on profiling repair dynamics over a time course – 4 h for (6-4)PP and 48 h for CPD. Genomic regions that harbor high-level early repair shortly after damage formation have not been studied, nor have their associated genomic and epigenomic characteristics been systematically explored.

Here, we performed XR-seq at times as early as 1 min for (6-4)PP and 12 min for CPD following UV irradiation to identify such repair hotspots. We systematically characterized the identified hotspots using additional high-throughput sequencing data that measures DNA damage formation, DNase I hypersensitivity, histone modifications, 3D chromatin interactions, and DNA replication timing. These extensive links between chromatin states and organization, cell cycle, DNA damage, and excision repair at specific genomic sites facilitate a better understanding of mutagenesis and carcinogenesis.

## RESULTS

### Genome-wide profiling of DNA excision repair kinetics through ordered high-throughput experiments

In this work, we present an experimental and analytical framework where we systematically assay DNA excision repair of UV-induced DNA damage over a time course. Figure 1A gives an outline of the experimental design. Specifically, we adopted XR-seq (7) to measure repair of (6-4)PPs at 1 min, 2 min, 5 min, 20 min, 1 h, 2 h, and 4 h and CPDs at 12 min in normal human skin fibroblasts (NHF1) after 20J/m_2_ UV treatment. The (6-4)PP XR-seq experiments from 1 min to 4 h were all performed with two biological replicates, with correlation coefficients greater than 0.99 between each pair of replicates (Figure S1). Quality control procedures show that all samples have good data quality with expected read length distributions and enrichments of TT and/or TC dinucleotides at the damage sites (Figure S2). To investigate the interplay between DNA damage and repair, we additionally adopted Damage-seq data (15) to profile the two types of UV-induced damage at 0 timepoint after UV irradiation. Refer to the Methods section for details on library preparation, sequencing, and bioinformatic analysis and Table S1 for detailed sample information.

**Figure 1.**
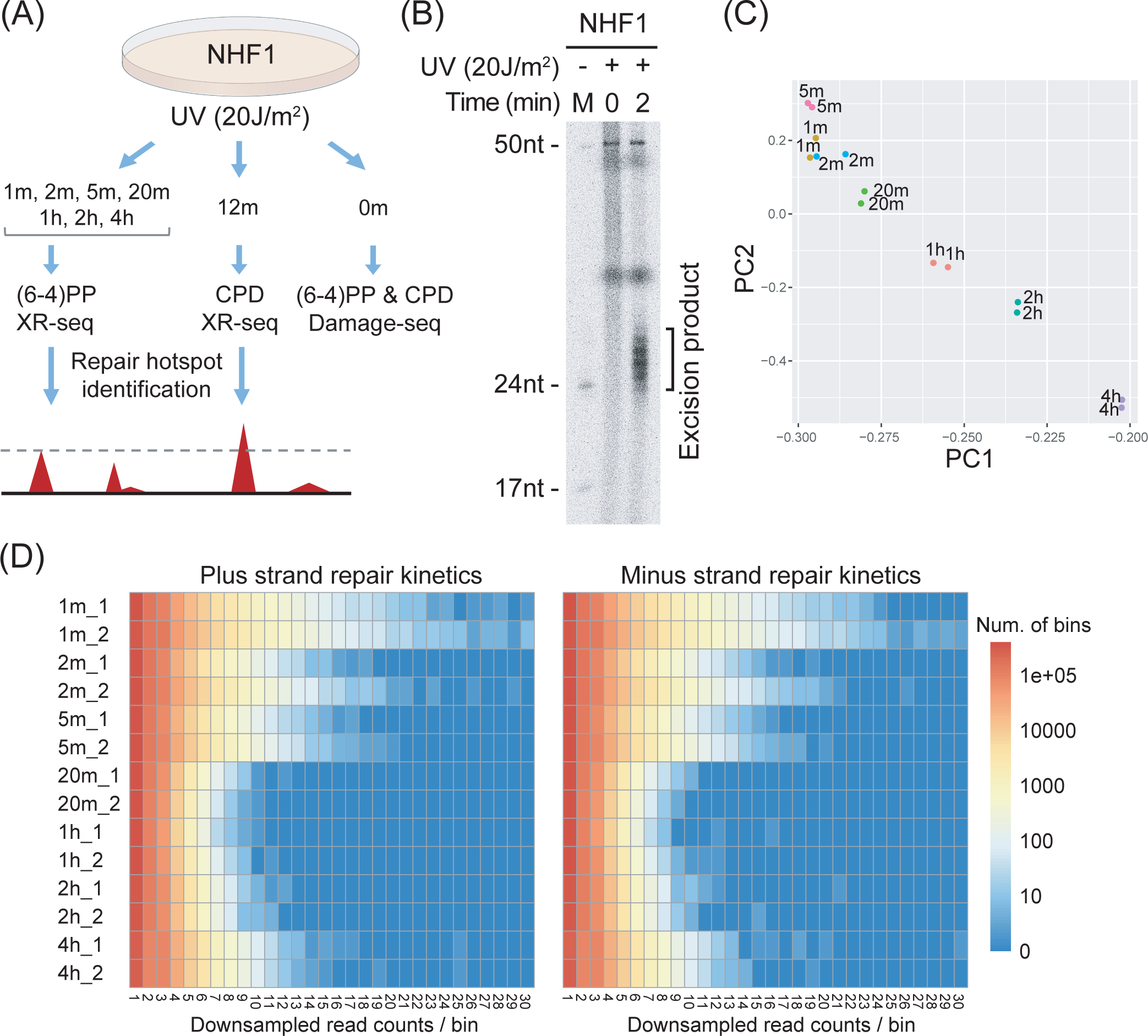
Excision repair kinetics and early-repair hotspots in normal human fibroblasts. (A) Experimental design to measure DNA damage and excision repair of the UV-induced (6-4)PP and CPD across different timepoints. (B) Detection of excision products at 0 min and 2 min timepoints *in vivo*. Following UV irradiation, the excised oligonucleotides were purified by TFIIH immunoprecipitation, radiolabeled, and resolved in a 10% sequencing gel. DNA excision products of (6-4)PP can be detected as early as 2 min upon damage induction. (C) Principal component analysis of genome-wide excision repair as measured by XR-seq shows repair kinetics across different timepoints. Between 1 min and 5 min, excised oligonucleotides were not degraded, and thus XR-seq measured cumulative repair. (D) Each row is a sample, and each column is a specific total number of reads per genomic bin. The color in the heatmap corresponds to the log counts of the number of bins with specific read depths. Early-repair hotspots exist in samples collected at early timepoints.

The bulk repair pathways and kinetics of these two types of DNA damage are known to be quite different (23,24). The global repair efficiently removes (6-4)PPs, with most of the damage excised within 4 h of UV treatment. We provide additional empirical evidence that the repair levels of (6-4)PP from the transcribed strand (TS) and the non-transcribed strand (NTS) are on par across all timepoints (Figure S3A-B). In contrast, CPDs are recognized primarily in a transcription-coupled manner, and their complete repair requires 12 to 48 h, depending on the UV dose. In our experiment, however, given the sampling timepoint as early as 12 min for CPDs, transcription-coupled repair has not contributed much, if at all, to the repair of CPDs. Specifically, as shown in Figure S3C, we do not observe a difference between TS and NTS repair at 12 min, which is likely too early for enough polymerases to have stalled and recruited the repair factors (25). This is of key importance so that our analysis is unbiasedly focused on the global repair of (6-4)PP at any timepoints and of CPD at the early 12 min timepoint, without transcription as a confounder. Refer to the Discussion section for more details.

In this study, we focus on characterizing global repair kinetics and identifying early-repair hotspots and late-repair coldspots. *In vivo* excision assay (24) by gel autoradiography at 0 min and 2 min reveals that the primary excision repair products can be detected as early as 2 min after UV treatment (Figure 1B). Using genome-wide repair data by XR-seq, we performed principal component analysis (PCA) (26) on the top 2,000 highly variable genes to generate a low-dimensional representation of the (6-4)PP repair data (Figure 1C). Since transcription-coupled repair does not contribute to the repair of (6-4)PPs, the PCA plots do not differ between the TS and NTS repair, except for reverted signs of the eigenvectors. Importantly, a repair trajectory on the principal component space can be reconstructed, which lines up well with the timepoints, suggesting differential repair kinetics over the time course (Figure 1C).

### Identification of early-repair hotspots and late-repair coldspots in normal human fibroblasts

We developed a computational framework to identify genome-wide early-repair hotspots using time-course XR-seq data. Briefly, we segmented the genome into consecutive bins of 50 bp long and identified bins that showed significantly enriched repair at earlier or later timepoints using a thresholding approach on the downsampled reads (Figure S4). We define such genomic bins as repair hotspots and coldspots, respectively. While such a method is efficient and effective in identifying the top few hundred repair hotspots and coldspots, we additionally adopted a more rigorous Poisson log linear model (27,28) on the read count data for normalization and testing of repair enrichment. The identified repair hotspots and coldspots show enriched repair levels compared to those as expected under the null (Figure S5). Refer to the Methods section for details.

Figure 1D shows the distributions of read counts per genomic bin across all (6-4)PP samples, and we note enrichment of both early repair at 1 min and late repair at 4 h, corresponding to repair hotspots and coldspots, respectively. For repair of (6-4)PPs, we identified on the genome-wide scale 175 and 156 repair hotspots from the plus and minus strand (Table S2) and 48 and 57 repair coldspots from the plus and minus strand (Table S3), respectively. For repair of CPDs, we focused on repair hotspots at an early timepoint to negate the global impact of transcription, and identified 99 and 93 repair hotspots from the plus and minus strands, respectively (Table S4). We could not search for intrinsic CPD coldspots, because 30 min after UV irradiation transcription-coupled repair becomes an important contributor to the repair profile.

The repair hotspots and coldspots reported are scattered across the entire human genome (Figure S6). XR-seq signals from examples of repair hotspots and coldspots are separated by strand and plotted across all timepoints in Figure 2. We also include epigenomic signals by DNase-seq, ChIP-seq from ENCODE (29), and Damage-seq signals at 0 min after UV treatment (15). Specifically, the XR-seq signals from an example (6-4)PP repair hotspot decrease dramatically from 1 min to 20 min and can be barely seen at 1 h (Figure 2A). In contrast, the XR-seq signals at a late-repair coldspot shown in Figure 2B increases over the time course and peaks at 4 h. Another representative CPD repair hotspot at 12 min is shown in Figure 2C.

**Figure 2.**
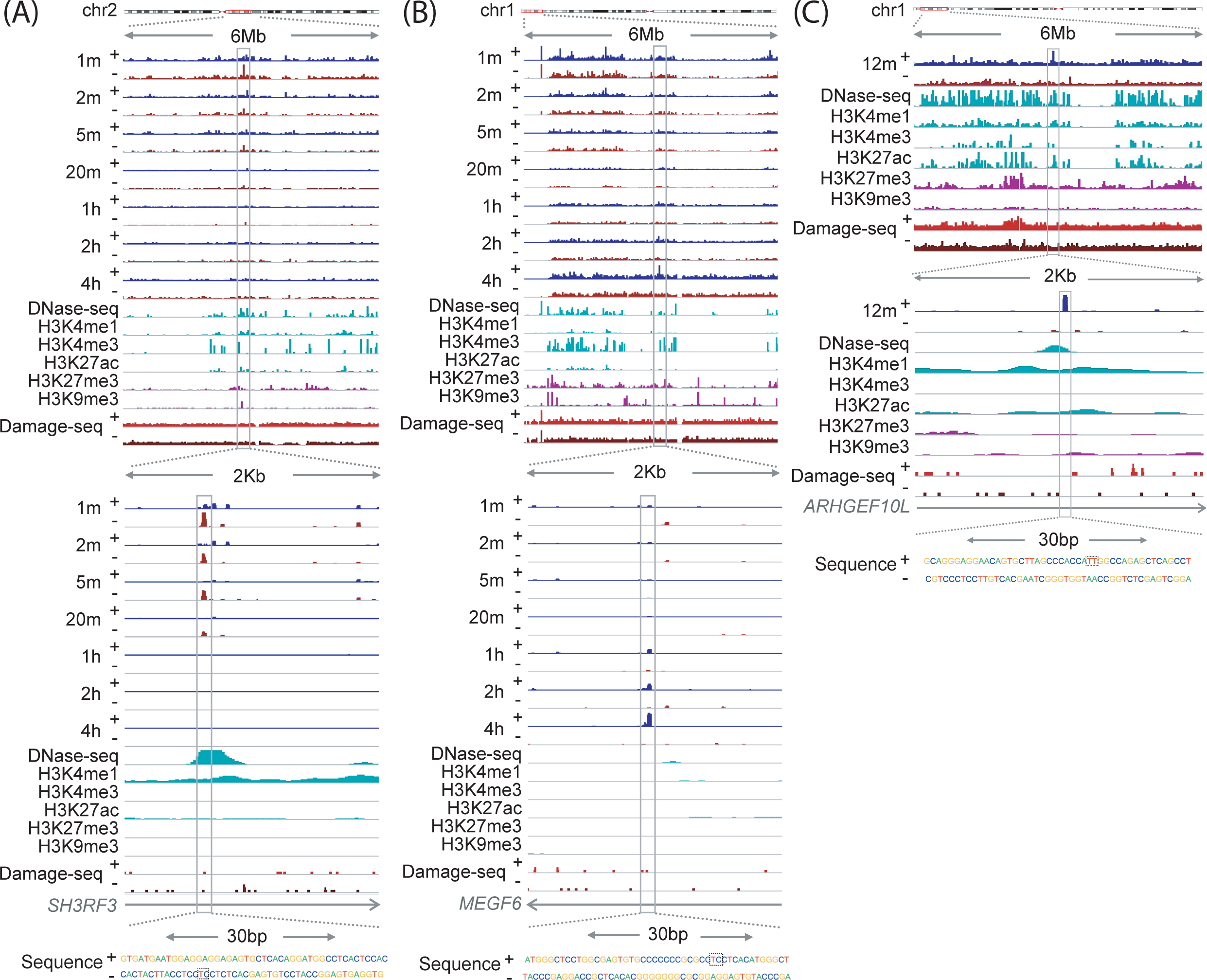
Distribution of DNA damage, repair, and epigenomic markers at identified repair hotspots and coldspots. XR-seq and Damage-seq data are shown for both strands, marked with + and -. Epigenetic data from ChIP-seq of histone modifications and DNase-seq are plotted on the same scale for cross comparison. Read count data normalized by sequencing depth are visualized in the Integrative Genomics Viewer. (A) A (6-4)PP repair hotspot from chr2. (B) A (6-4)PP repair coldspot from chr1. (C) A CPD repair hotspot from chr1. All examples are from intronic gene regions overlapping annotated enhancers. Zoomed-in view of canonical sequences is overlaid in the bottom, with the damage/repair sites shown in dashed boxes.

We next performed sequence context analysis using all reads that are mapped to the repair hotspots and coldspots, respectively. We trimmed the reads to be 15 bp long centering at the damage sites and calculated strand-specific nucleotide frequencies in repair hotspots, coldspots, and randomly picked spots. Interestingly, we identified an enrichment of cytosine in the flanking regions of the damage/repair sites for both repair hotspots and coldspots (Figure S7). Motif analysis by the MEME suite (30) confirmed the enrichment of cytosine adjacent to the repair sites, which are themselves enriched with canonical sequences of CTCA for (6-4)PP and TT for CPD (Table S5). While it has been previously shown that active transcription factor binding sites have decreased levels of excision repair (21), here we report a preference for cytosine at both early and late timepoints, which have increased levels of repair compared to null regions. Notably, such cytosine enrichment in the flanking regions of the repair sites is not due to sequence context bias, as we show in Figure S8 that not all genomic regions enriched for cytosines are enriched for excision repair. While an interesting observation, the underlying biological mechanisms need further investigation.

### Early-repair hotspots are not hypersensitive to DNA damage

A recent study assayed DNA damage by next-generation sequencing and reported a total of 153 hyper-hotspots acquiring CPDs much more frequently than the genomic average in primary human fibroblasts (31). Of these damage hotspots, 83 are from the plus strand and 74 are from the minus strand, each having at least five recurrent sequence reads (31). To investigate whether the identified repair hotspots simply result from increased levels of DNA damage, we first intersected the reported damage hotspots from Premi et al. (31) with the 99 and 93 CPD repair hotspots from the plus and minus strand that we identified. We found that none of our identified repair hotspots overlapped with the reported damage hotspots.

To further confirm and replicate this seemingly striking result, we analyzed genome-wide CPD DNA damage data generated by our previously developed Damage-seq protocol (15), to quantify damage levels at 0 timepoint after UV irradiation with single-nucleotide resolution. After stringent quality control procedures (refer to the Methods section for details), we identified 91 and 78 CPD damage hotspots from the plus and minus strand, respectively, each having at least ten mapped reads (Table S6). Notably, these Damage-seq hotspots are shown to be enriched for heterochromatin and repressed regions (Figure S9), which is concordant with previous reports (32,33). Importantly, none of the CPD damage hotspots, identified from this parallel Damage-seq platform, overlap with the repair hotspots.

In addition, we compared the DNA damage levels for (6-4)PP and CPD from three independent sequencing technologies – Damage-seq (15), adductSeq (31), and CPD-seq (19) – at our identified hotspots and coldspots against those from randomly sampled regions along the genome. To account for the sparse sampling when measuring DNA damage by next-generation sequencing, we also extend the regions corresponding to the repair hotspots, coldspots, and random spots at both ends for 20bp and 500bp, respectively. Our results, shown in Figure 3, suggest that there is no significant difference in the damage levels between the three repair categories (hotspot, coldspot, and random spot). The zoom-in and zoom-out views of three example repair hotspots and coldspots in Figure 2 also suggest that the Damage-seq reads are uniformly distributed in the flanking regions. Previous results have demonstrated that the UV-induced DNA damage is indeed virtually uniform across the entire human genome, while repair is affected by chromatin states, transcription factor binding, etc., in a manner dependent on the type of DNA damage (15). While we note that the shallow depth of coverage of Damage-seq can be a limiting factor (refer to the Discussion section for details), our results validate our conclusion that the identified repair hotspots are not damage hotspots.

**Figure 3.**
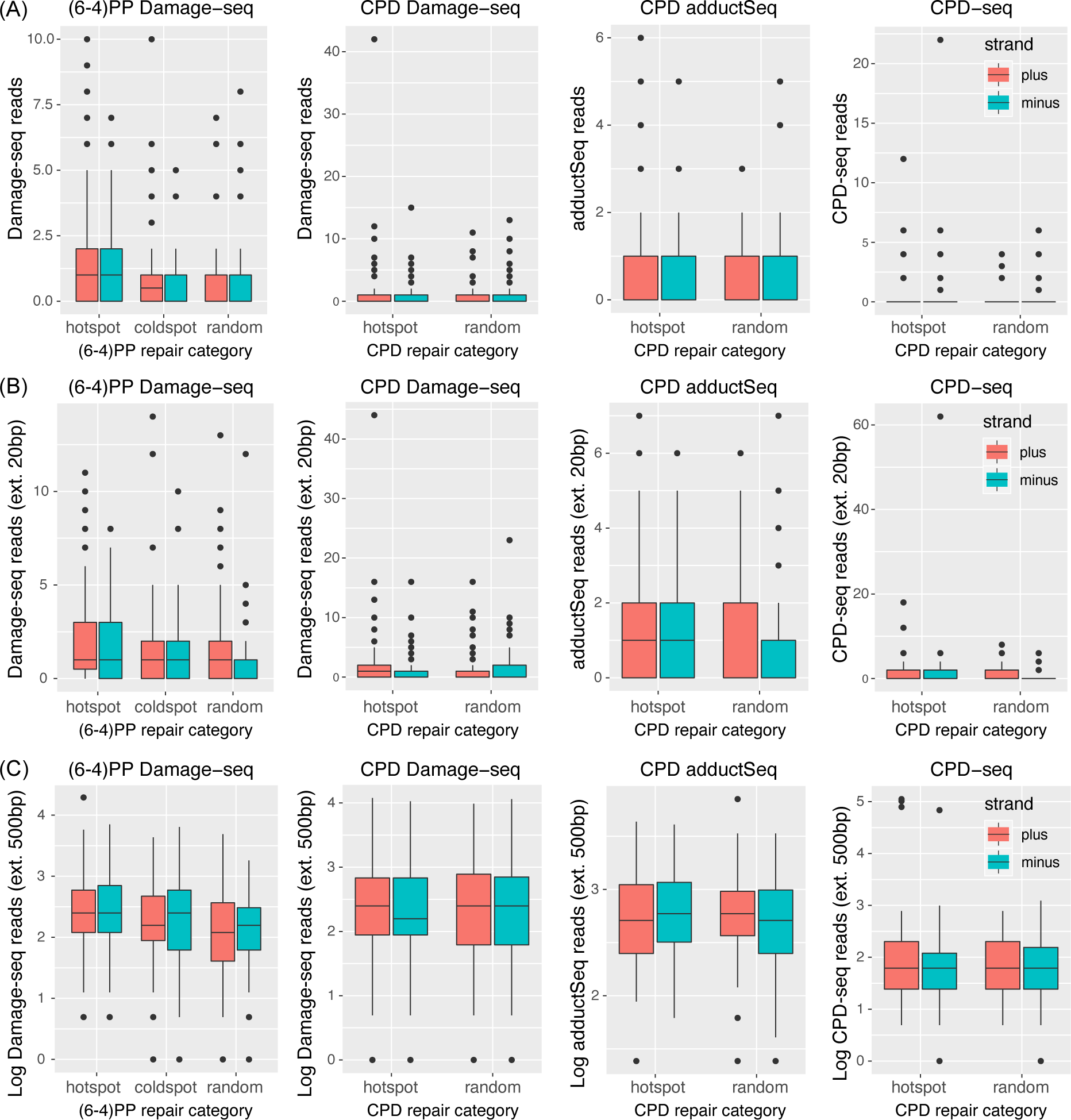
Repair hotspots are not hypersensitive to ultraviolet radiation. There is no enriched DNA damage, as measured by Damage-seq, adductSeq, and CPD-seq, at the identified repair hotspots for (6-4)PP and CPD. (A) Read counts for DNA damage are computed in repair hotspots, coldspots, and random spots. The regions corresponding to the different repair categories are extended at both ends for 20bp and 500bp, respectively, to account for the shallow sequencing depth by quantifying DNA damage.

### Early-repair hotspots are enriched for promoters and enhancers in open chromatin regions

The packaging of DNA into chromatin can hinder the access of repair proteins and affect the repair efficiency (16,17) and specific histone modifications are associated with different functional and cytological chromatin states. To investigate the relationship between the identified repair hotspots/coldspots and the epigenomic markers, we used publicly available DNase-seq data and histone modification ChIP-seq data for an adult human fibroblast cell line (NHLF) from ENCODE (29). Our results suggest that the earliest repair occurs preferentially in active and open-chromatin regions. Specifically, chromatin accessibility by DNase-seq is significantly higher in repair hotspots than in random genomic regions, and it is significantly lower in repair coldspots (Figure 4). For histone modifications, repair hotspots have significantly higher ChIP-seq signals compared to random spots for activation markers, including H3K4me1, H3K4me3, and H3K27ac (Figure 4).

**Figure 4.**
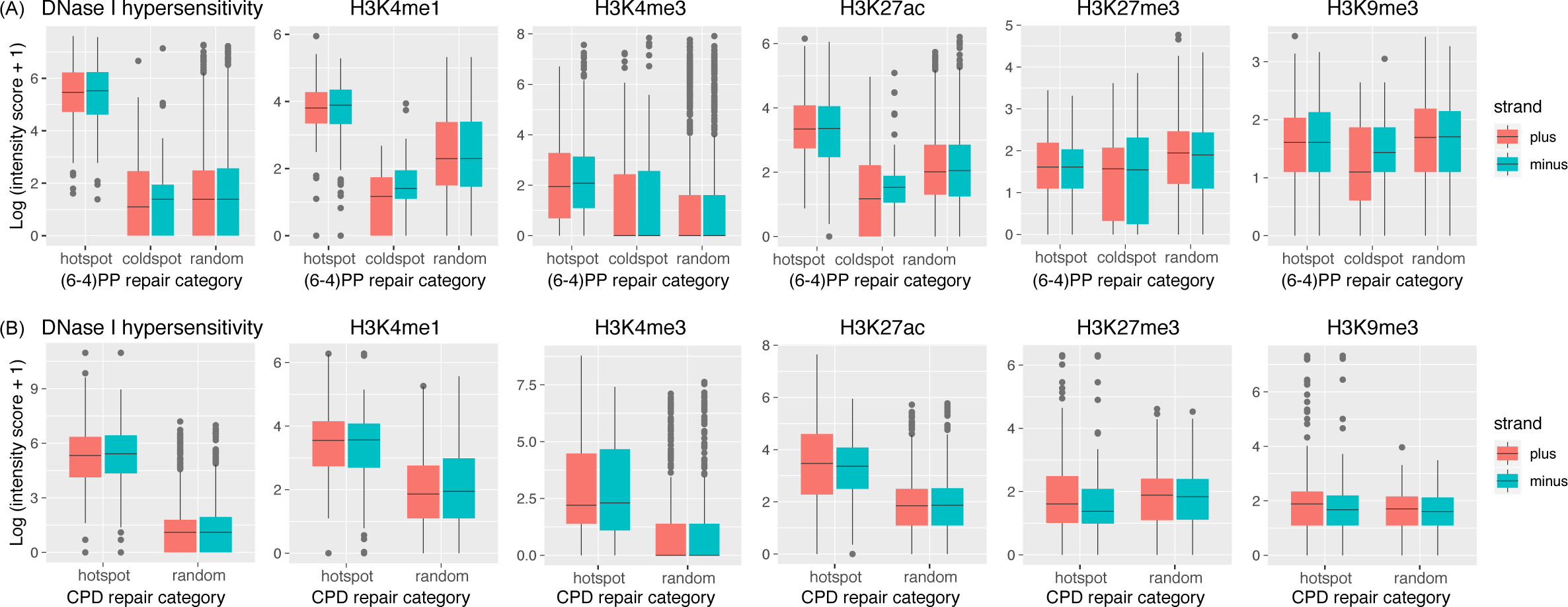
Genome-wide repair hotspots are associated with epigenomic markers. Chromatin accessibility (DNase I hypersensitivity) is higher for repair hotspots and lower for coldspots. Repair hotspots are also characterized by higher ChIP-seq signals for H3K4me1, H3K4me3, and H3K27ac, markers for gene activations.

We adopted the segmented chromatin states by chromHMM (34) to annotate the repair hotspots for (6-4)PP and CPD and found a significant enrichment of active promoters and enhancers (Figure 5A), which are characterized by nucleosome loss and open chromatin. Genome-wide repair kinetics inferred across timepoints confirm that enhancers are repaired at earlier timepoints and that repressed/heterochromatin regions are repaired at later timepoints (Figure S10). Additionally, we performed genome-wide annotations for CpG islands and genic/intergenic regions. Our results suggest that the repair hotspots are enriched for inter CpG regions (i.e., depleted for CpG islands) (Figure 5B) and intronic regions (Figure 5C), both of which are enriched for AT and presumably result in more repair. Notably, the number of identified repair coldspots is too low to generate reproducible annotation results between the replicates.

**Figure 5.**
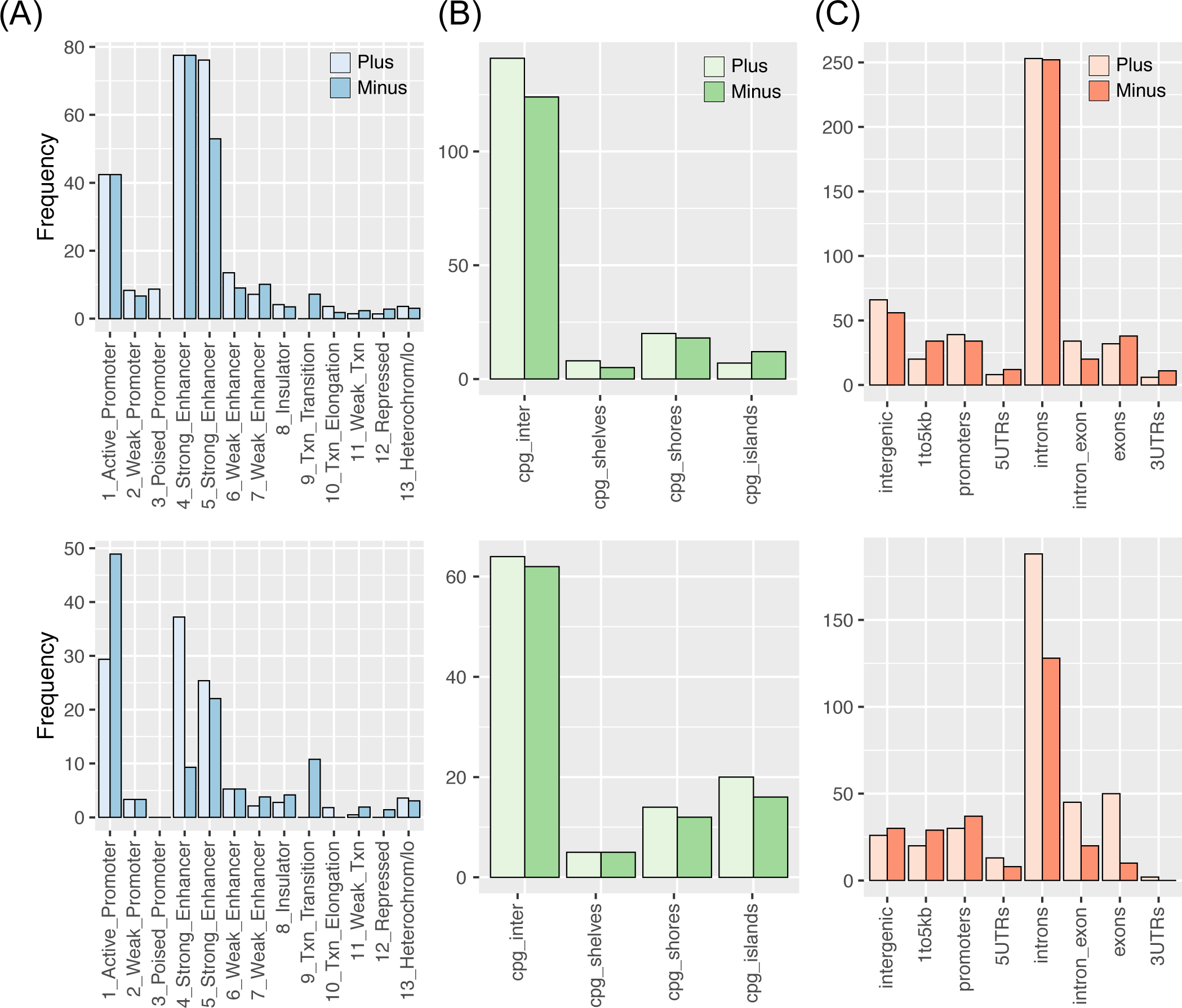
Genome-wide repair hotspots are enriched for enhancers. Annotations for (A) chromatin states, (B) CpG islands, and (C) genic/intergenic regions are shown for repair hotspots for (6-4)PP (top) and CPD (bottom), separated by strands. Repair hotspots are enriched for enhancers and promoters, which are in open-chromatin regions. Repair hotspots are also enriched in inter-CpG and intronic regions, both of which are AT-rich.

### Early-repair hotspots are enriched for frequently interacting regions (FIREs) and super-enhancers

With the genome-wide enrichment of active promoters and enhancers in open chromatin regions of the detected repair hotspots, we hypothesize that the 3D genome structure contributes to the observed differential repair kinetics between different regions of the genome. To test this hypothesis, we sought to interrogate the publicly available Hi-C data of human fibroblast cell line IMR90 (35,36). Specifically, after quality control procedures and data normalization, we profiled frequently interacting regions (FIREs) using FIREcaller (37). After overlapping the repair hotspots and coldspots with the called FIREs (Table S7A), we found that a significantly higher proportion of repair hotspots overlap with FIREs – 23.16% and 11.76% for (6-4)PP and CPD, respectively – compared to a genome average of 6.93% based on the profiled FIREs (Figure 6A). Conversely, the overlapping proportion of (6-4)PP repair coldspots is 3.23%, significantly lower than the genome average (Figure 6A).

**Figure 6.**
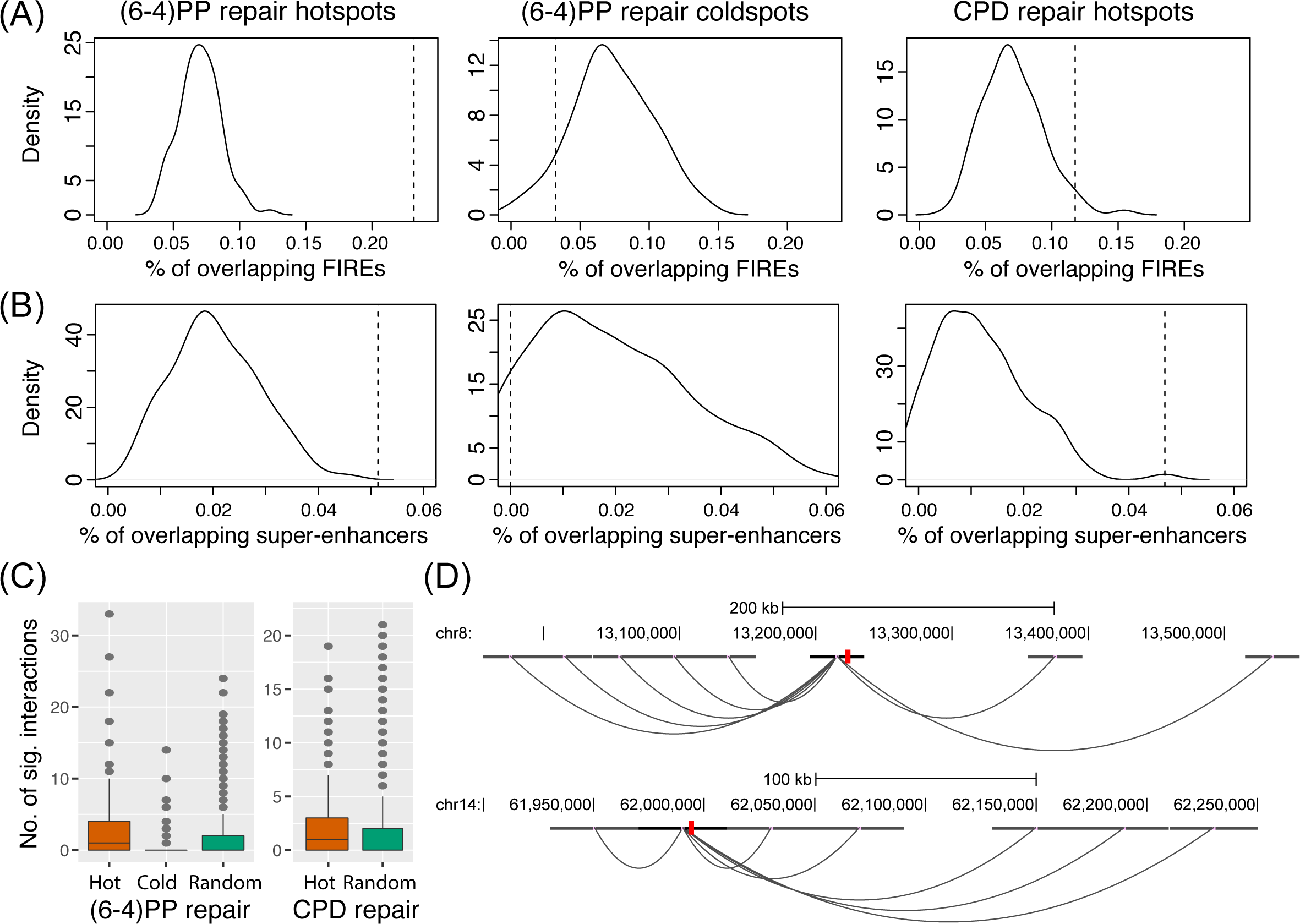
Genome-wide repair hotspots overlap with FIREs and super-enhancers identified by Hi-C. FIREs and super-enhancers were identified and annotated using Hi-C data from human fibroblasts. Repair hotspots overlap with (A) FIREs and (B) super-enhancers with significantly higher proportions compared to the genome-wide averages. The solid density curves are generated from bootstrapping different regions along the genome as the null case; the dashed vertical lines are the observed proportions for the repair hotspots and coldspots. (C) Repair hotspots have a significantly higher number of significant interactions, identified by Hi-C. (D) Two examples of (6-4)PP repair hotspots (chr8:13224201-13224300 and chr14:61994601-61994700) that overlap with both FIREs and super-enhancers, and loop to different regions of the genome. Hi-C data has low resolution, and the significant interactions are drawn from the center of each bin, which does not exactly overlap with the identified hotspot shown in red.

FIREs have been previously reported to be enriched for super-enhancers (38). We have demonstrated that the repair hotspots are enriched for both FIREs and enhancers. We also observed that, across many cases, multiple enhancers that overlapped with the repair hotspots were from the same genomic regions (Figure S11). As such, we hypothesize that the repair hotspots are also enriched for super-enhancers and thus additionally adopted a list of previously annotated super-enhancers in the human fibroblasts (39) (Table S7B). We found that, compared to a genome-wide average of 2.05%, the early-repair hotspots are indeed enriched for super-enhancers (5.14% and 4.69% for (6-4)PP and CPD repair hotspots, respectively), while none of the repair coldspots overlap with super-enhancers (0% for (6-4)PP repair coldspot) (Figure 6B).

In addition, we called significant interactions based on the Hi-C contact matrix using the Fit-Hi-C method (40) (Table S7C) and showed that early-repair hotspots also overlap with a significantly higher number of significant interactions (Figure 6C). Refer to the Methods section for details on data analysis. The overlapping information of the called repair hotspots and coldspots with the profiled FIREs, super-enhancers, and significant chromatin interactions are included in Table S8. Figure 6D illustrates the loop interactions of two identified repair hotspots. Notably, these two hotspots also overlap with both FIREs and super-enhancers. These results collectively provide a global picture of genetic regulation of repair kinetics via 3D genome structures.

### Early-repair hotspots are enriched for early-replication domains

We next set out to test the relationship between the repair kinetics and replication timing. We used the genome-wide Repli-Seq data of the human fibroblast cell line IMR90, which maps high-resolution DNA replication patterns with respect to both cell-cycle time and genomic position (41). A recent re-analysis of the data using a deep-learning model (42) segmented the genome into different replication domains, including the early-replication domain (ERD), late-replication domain (LRD), up-transition zone (UTZ), and down-transition zone (DTZ). Refer to Table S9 for genome-wide segmentation results. When overlapping the identified repair hotspots with the segmented replication domains, we find that early-repair hotspots are also significantly enriched for early-replication domains (Figure 7). While replication time has been shown to be correlated with chromatin accessibility (41), here we have demonstrated a potential cell-cycle effect on DNA excision repair. That is, regions that are duplicated early also tend to be repaired early, and this has been shown to result in mutagenesis asymmetries (17) – we further elaborate such implications in the Discussion section.

**Figure 7.**
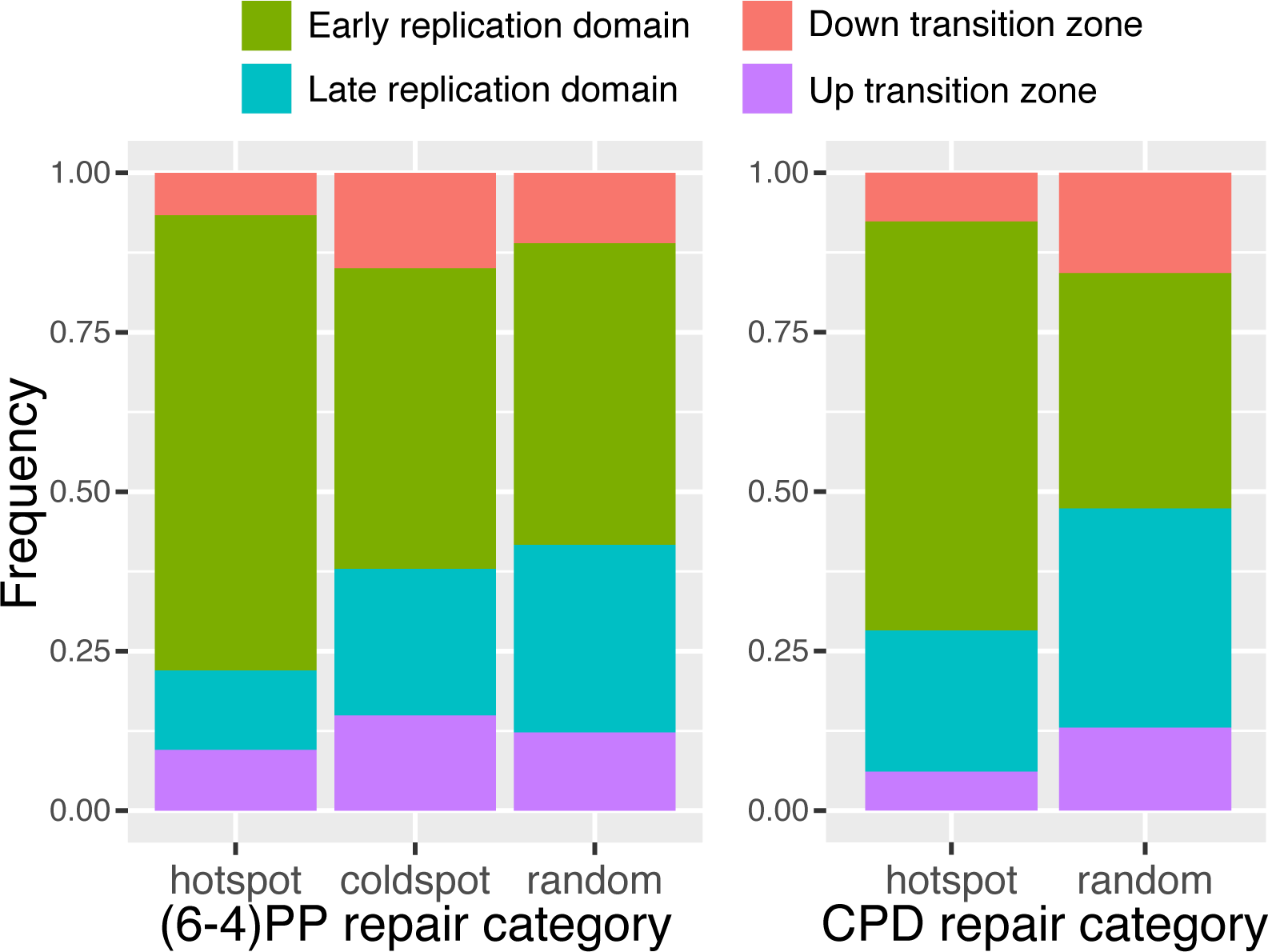
Genome-wide repair hotspots are enriched for early-replication domains. Replication timing domains were identified using Repli-Seq data. Compared to genome-wide average (random), there is a significantly higher proportion of the early-repair hotspots located in the early-replication domains.

### Profiling gene-level repair kinetics in normal human fibroblasts

So far, our analysis has been conducted on the whole-genome level. Identifying dynamic changes at individual gene levels can provide important insights about genes and pathways that exhibit significant changes in repair dynamics (43). We calculated strand-specific reads per kilobase per million reads (RPKM) for all the genes in the human genome. RPKMs across nine genes associated with circadian rhythm (e.g., *CLOCK, BMAL1, PER1, CRY1*) and excision repair (e.g., *XPA, XPC, CSA, CSB*) present different repair dynamics over the time course (Figure S12). We adopted Trendy (44) to perform segmented regression analysis (Table S10) on the ordered XR-seq data. We focused on significant genes with adjusted *R*2 greater than 0.8 from the regression analysis and identified 3,017 significant genes with fitted trend as “down,” 3,496 significant genes with fitted trend as “up-down,” 1,024 significant genes with fitted trend as “up,” and 1,452 significant genes with fitted trend as “down-up” (Figure S13). The number of genes that show the “down” trend is two-fold higher than the number of genes that show the “up” trend, which is consistent with our previous observation – there are more repair hotspots than coldspots. For these four groups of significant genes, we further performed gene ontology (GO) enrichment analysis using PANTHER (45). For genes with the “down” and “up-down” trend, GO biological processes are enriched for a large number of biological regulations, including regulation of the metabolic process, gene expression, developmental process, cell cycle, and cell death, etc. (Table S11A-B). On the other hand, there are no obvious GO enrichment terms for the “up” or “down-up” genes (Table S11C-D).

## DISCUSSION

We adopted time-course XR-seq data to detect genome-wide repair hotspots and coldspots and integrated additional omics data from various next-generation sequencing platforms to investigate the relationship between DNA damage, excision repair, epigenomic markers, 3D genome organization, and replication timing. To our best knowledge, this, for the first time, demonstrates the connection between repair kinetics and chromatin organization (46) with high resolution in human cell lines. The identified repair hotspots can serve as additional functional genomic features in studies of the human genome at the multidimensional level. We believe the framework and multi-omics data we present here can be useful to scientists both in the fields of chromatin dynamics and those interested in determinants of repair rates in the human genome. Our approach is applicable to studying the response to other types of DNA damage, including that induced by drugs used in chemotherapy and thus has the potential for informing new therapeutic strategies.

Using paired XR-seq and Damage-seq data, we provided additional empirical evidence that the damage levels in the regions of the repair hotspots were not enriched, in concordance with previous reports (14,15). Therefore, we attributed the observed differences in repair patterns to heterogeneous repair efficiencies and not to damage formation. It is, however, noteworthy that the depth of coverage by Damage-seq is an order of magnitude lower than that by XR-seq. Specifically, the genome-wide average total number of reads between the two replicates for (6-4)PP Damage-seq is 20.5 million, while that average for (6-4)PP XR-seq is 181.8 million (Table S1). As such, it is possible that we do not have enough sequencing depth to detect Damage-seq hotspots with high sensitivity. The fact that a good proportion of Damage-seq reads get mapped to heterochromatin regions further complicates analysis and lowers detection power. This issue also persists for the recently developed adductSeq and FreqSeq, where a Poisson model was adopted with a genomic average rate of 0.07 reads per pyrimidine dinucleotide in fibroblasts (31). A Poisson distribution with a mean parameter under the null much less than one could potentially suffer from overdispersion and inflated false positive rates. Targeted Damage-seq offers a solution for both discovery and validation. However, as with targeted DNA sequencing, the data can be extremely noisy due to targeting, amplification, and sequencing biases and artifacts (47).

Using the genome-wide profile of DNA replication timing, we demonstrate that the early-repair hotspots also tend to reside in early-replication domains. The inevitable effect of replication on gene dosage and copy number could also have regulatory consequences (48). Genes/regions that are duplicated early will be present at twice the copy number of late replicating domains for most of the duration of S phase, increasing both the DNA amount (as sister chromatids) and the transcriptional output. The ability to perform excision repair for a greater fraction of the cell cycle has been shown to result in lower mutation rates in the early-replication domains (49,50).

UV-induced DNA damage and excision repair have been linked with mutagenesis and carcinogenesis. Recent studies have shown that mutation hotspots exhibit strong increases in CPD formation efficacy (51-53) and that excision repair is attenuated in transcription factor binding sites, leading to elevated somatic mutation rates (19,21). In terms of repair kinetics, Adar et al. (17) showed that globally late-repaired regions are associated with a higher level of cancer-linked mutations. To further investigate this in our study context, we quantified the rates of somatic point mutations associated with melanoma (1) in the repair hotspots and coldspots. However, due to the low number of hotspots and coldspots, as well as their small genomic lengths, we found very few overlapped with mutations and were thus underpowered to test for the differences in mutation rates between different repair categories.

While the repair of (6-4)PP is predominantly carried out by global repair, transcription-coupled repair plays an essential role in removing the more abundant and less helix-distorting CPD adducts. In this study, we have intentionally focused on profiling global repair hotspots for both (6-4)PP and CPD at early timepoints without the effect of transcription-coupled repair. Genes exhibit both transcriptional dynamics over a time course and biological fluctuations due to transcriptional bursting (54) at the same timepoint. This confounds and complicates the analysis. To disentangle the effects of global repair and transcriptional-coupled repair in identifying repair hotspots is an unsolved yet challenging problem.

## METHODS

### Experimental methods

#### Cell culture and UV irradiation

Human NHF1 cells were obtained from W.K. Kaufmann (University of North Carolina, Chapel Hill) (55) and cultured in Dulbecco’s Modified Eagle Medium (DMEM) with 10% FBS at 37 °C in a 5% CO_2_ humidified chamber. For (6-4)PP XR-seq at 1 min and 2 min timepoints, UV irradiation was performed as previously described (7,56). Briefly, the 80% confluent NHF1 cells in one petri dish were irradiated for 20 s under a 250 nm UV lamp (1 J/m_2_/s) after removing the culture medium. 37 °C DMEM with 10% FBS medium was immediately added into the petri dish, then the medium was poured off and the petri dish was put on ice promptly at the end of 1 min or 2 min after UV irradiation. The time count starts from the end of 20 s UV irradiation and ends at the timepoint when the petri dish is put on ice. The cells were washed one time with ice-cold PBS before being harvested by a cell scraper in 10 ml ice-cold PBS. In each replicate of (6-4)PP XR-seq experiment, 50 and 30 petri dishes (150mm x 15mm) containing NHF1 cells were treated one by one at 1 min and 2 min timepoints, respectively. Cell culture, UV treatment, and library preparation for (6-4)PP XR-seq at 5 min, 20 min, 1 h, 2 h, and 4 h, and CPD XR-seq at 12 min were performed in previous studies (7,10,17). For *in vivo* excision assay, UV irradiation was performed as aforementioned, and 10 and 5 petri dishes (150mm ⨯ 15mm) containing NHF1 cells were used at 0 min and 2 min timepoints respectively.

#### Excision assay

The *in vivo* excision assay was performed as described (24,57). Following UV irradiation, the excision products were isolated by gentle cell lysis and nonchromatin fraction separation and purified by TFIIH immunoprecipitation. The purified excision products were then 3’ radiolabeled by terminal deoxynucleotidyl transferase and [α-_32_P]-3’-dATP, and resolved in a 10% denaturing acrylamide gel. Ten and five petri dishes (150mm x 15mm) of NHF1 cells were used at 0 min and 2 min, respectively.

#### XR-seq library preparation and sequencing

XR-seq libraries were prepared as described in the previous protocol (57). Briefly, the excision products were isolated by TFIIH immunoprecipitation following gentle cell lysis and non-chromatin fraction separation, and ligated with adaptors. The ligated excision products were then further purified by immunoprecipitation with anti-(6-4)PP antibody and repaired by (6-4)PP photolyase before the library amplification by PCR. Libraries were sequenced on an Illumina HiSeq 4000 platform.

### Data collection

(6-4)PP XR-seq data at 5 min, 20 min, 1 h, 2 h, and 4 h were downloaded from the Gene Expression Omnibus (GEO) with accession numbers GSE67941 (7) and GSE76391 (17). CPD XR-seq data at 12 min were downloaded from GEO with accession number GSE138846 (10). CPD and (6-4)PP damage data of NHF1 by Damage-seq were downloaded from GEO with accession number GSE98025 (15); CPD damage data of NHF1 by CPD-seq were downloaded from GEO with accession number GSM2772322 and GSM2772323 (19); CPD damage data of human primary fibroblast by adductSeq were downloaded from GEO with accession number GSM4073616 and GSM4073634 (31). Hyper-hotspots for UV-induced CPD damage in primary human fibroblasts were downloaded from Premi et al. (31). NHDF H3K4me1 (ENCODE Data Coordination Center accession number ENCSR000ARV), H3K4me3 (accession number ENCSR000DPR), H3K27ac (accession number ENCSR000APN), H3K27me3 (accession number ENCSR000APO), H3K9me3 (accession number ENCSR000ARX), and DNase-seq (accession number ENCSR000EMP) data were downloaded from the ENCODE portal (29). NHLF chromatin state segmentation results by chromHMM were downloaded from UCSC accession number wgEncodeEH000792 (34). Hi-C data of IMR90 were downloaded from GEO with accession number GSE43070 (35) and from https://bioconductor.org/packages/HiCDataHumanIMR90/ (36). The list of annotated super-enhancers in IMR90 was downloaded from the Roadmap Epigenomics Consortium (39). Genomic categories of replication timing from Repli-Seq data of IMR90 (41) were downloaded from GSE53984 (42).

### Bioinformatic and statistical analysis

#### XR-seq bioinformatic pre-processing

For XR-seq, cutadapt (58) was used to trim reads with adaptor sequence TGGAATTCTCGGGTGCCAAGGAACTCCAGTNNNNNNACGATCTCGTATGCCGTCT TCTGCTTG at the 3’ end and discard untrimmed reads. BWA (59) was used for alignment of single-end short reads. Unmapped reads and reads that map to multiple locations with the same alignment quality were removed using Samtools (60). Post-alignment filtering steps were adopted using Rsamtools (http://bioconductor.org/packages/Rsamtools/). Specifically, if multiple reads share the same 5’ and 3’ end coordinates, we keep only one to perform deduplication. We also only keep reads that have mapping quality greater than 20 and are of lengths 21 bp to 31 bp.

#### Gene-level quantification of excision repair

Reads from the TS and NTS strands were separated using known gene annotations for hg19 by ENSEMBL. We use RPKM for within-sample normalization for the XR-seq data. To perform gene-level quantification and downstream analysis including segmented regression, we adopted a stringent quality control procedure and only retained genes that: (i) had at least ten TT or TC dinucleotides from either TS or NTS; (ii) were less than 300Kb; and (iii) had at least ten reads in total across all XR-seq samples. In addition, we took the ratio of the reads from the TS and the NTS [TS/(TS+NTS)] to remove biases and artifact that are shared between the two DNA strands, i.e., library size, gene length, and other gene-specific biases, such as sequencing bias and antibody pulldown efficiency, etc. The ratio is bound between 0 and 1 and sheds light upon how transcription-coupled repair and global repair interplay (Figure S3).

#### Identification of repair hotspots and coldspots

We started by segmenting the human reference genome into consecutive bins of 50 bp long. We then calculated the observed depth of coverage per bin by XR-seq, separating the plus-strand reads (+) and the minus-strand reads (-). To mitigate the effect of library size/sequencing depth, we downsampled the reads in each sample to 7.7 million without replacement. To identify repair hotspots and coldspot, we set a threshold on the number of read counts per genomic bin in the 1 min and 4 h samples. Specifically, to identify (6-4)PP repair hotspots, we require at least 15 reads mapped in both replicates at 1min and at most 5 reads mapped in both replicates at 4 h. The read count threshold is relaxed for the identification of coldspots, which have a smaller number compared to the hotspots. For CPD repair, to avoid complicatedness due to transcription-coupled repair at later timepoints, we focused on CPD repair hotspots only.

In addition to the thresholding approach, we adopted a more rigorous cross-sample Poisson log linear model (27,28) for data normalization. Specifically, we denote *Y* as the observed repair matrix, with row *i* corresponding to the *i*th genomic bin and column *j* corresponding to the *j*th sample. The “null” model, which reflects the expected coverage when there is no biologically relevant repair enrichment, is

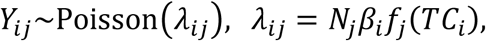

where *N*_*j*_ is the total number of mapped reads for sample *j*(fixed for downsampled data), β_*i*_reflects the bin-specific bias due to library preparation and sequencing bias, and *f*_*j*_(*TC*_*i*_) is the sample-specific bias due to TC (thymine and cytosine) content for damage/repair. The goal of fitting the null model to the data is to estimate the various sources of biases, which can then be used for normalization. We adopt a robust iterative maximum-likelihood algorithm (28) for estimating the parameters of the null model. Plus and minus strands are analyzed separately.

Given a first-pass of the calling algorithms, we identified strong repair hotspots in pericentromeric regions, which were collapsed repeats annotated as unique sequences in the reference genome (e.g., ribosomal DNA (9)). It is important to exclude artifacts as stringently as possible, and thus we undertook an additional quality control step. “Blacklist” bins, including segmental duplication regions (http://humanparalogy.gs.washington.edu/build37/data/GRCh37GenomicSuperDup.tab), gaps in reference assembly from telomere, centromere, and/or heterochromatin regions (https://gist.github.com/leipzig/6123703), and repeating elements by RepeatMasker (https://genome.ucsc.edu/cgi-bin/hgTrackUi?g=rmsk) are masked in downstream analysis.

#### Hi-C data analysis

We adopted the Hi-C data of human fibroblast cell line IMR90 (35,36) to investigate the relationship between identified repair hotspots and the 3D genome structure. We took the raw contact matrix with 40 kb resolution as input and detected FIREs, which play important roles in transcriptional regulations, across the entire genome using FIREcaller (37). To further investigate whether these repair hotspots are involved in functional chromatin looping between regulatory elements and their target genes, we adopted the Fit-Hi-C approach (40) to identify long-range chromatin interactions on all 40 kb bin pairs within a maximal 3 MB region. The interactions with *p*-value < 2.31e-11 were considered as statistically significant (61).

## DATA AND CODE AVAILABILITY

The data reported in this paper have been deposited in GEO with accession number GSE148303. Scripts used in this paper are available at https://github.com/yuchaojiang/damage_repair.

## ACKNOWLEDGEMENTS

This work was supported by NIH Grants R35 GM118102 (to A.S.), R01 ES027255 (to A.S.), R35 GM138342 (to Y.J.), K99 ES030015 (to W.L.), and a pilot award from the UNC Computational Medicine Program (to Y.J.). We thank the Sancar Lab members and Drs. Sheera Adar, Ming Hu, and Jeremy Simon for useful comments and feedback.

## AUTHOR CONTRIBUTIONS

AS envisioned and initiated the study, while LW and LAL-B performed the experiments. YJ devised the analytical framework, and all authors executed the data analysis. YJ wrote the manuscript, with contributions from WL on experimental methods. The manuscript was further edited and approved by all authors.

