## Supplementary Materials for "Nucleotide excision repair hotspots and coldspots of UV-induced DNA damage in the human genome"

**Figure S1: XR-seq signals between two biological replicates.** (6-4)PP DNA repair levels between each pair of biological replicates collected at (A) 1 min, (B) 2 min, (C) 5 min, (D) 20 min, (E) 1 h, (F) 2 h, and (G) 4 h. The two (6-4)PP XR-seq replicates across all measured timepoints have correlation coefficients above 0.99, indicating good data quality and reproducibility.

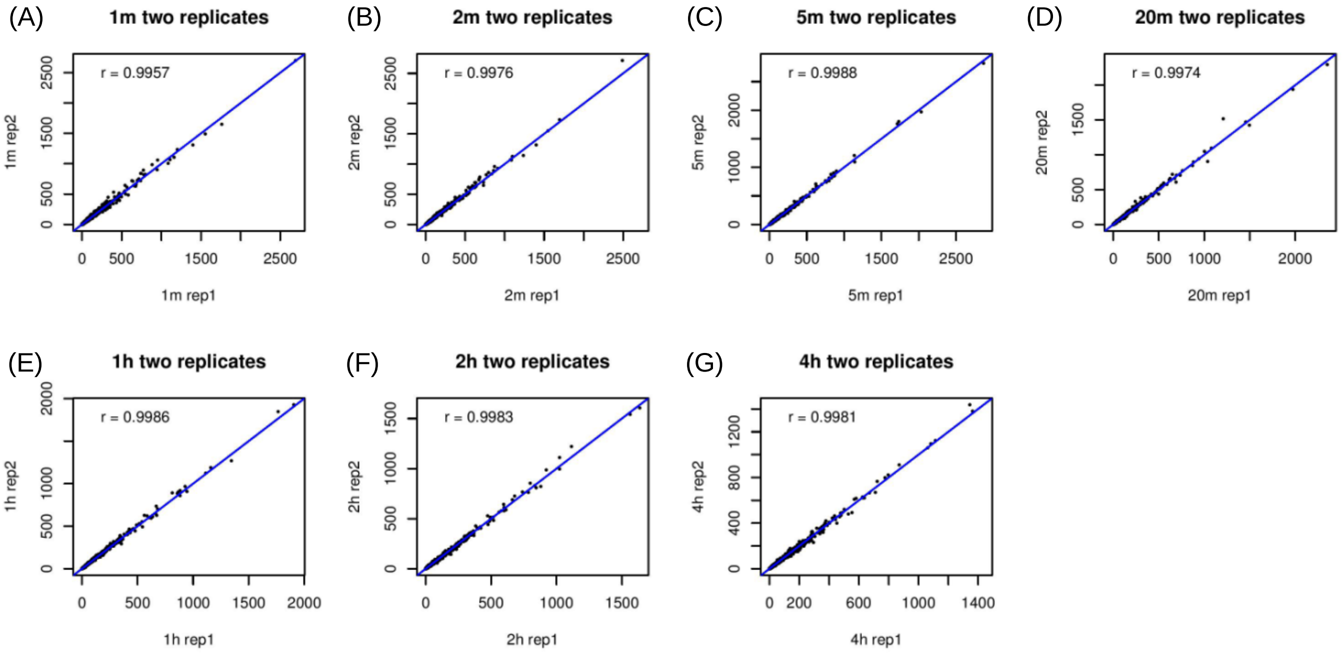

**Figure S2: Read length distribution and di-nucleotide frequency of XR-seq data.** Lengths of the excised oligomers fall within the range of 21 to 31 bp across all samples with peak at 26-27nt; TT/TC dinucleotides are enriched at the damage/repair sites by XR-seq. Three samples from early and late timepoints are shown: (A) (6-4)PP XR-seq 1 min replicate 1; (B) (6-4)PP XR-seq 4 h replicate 1; and (C) CPD XR-seq 12 min.

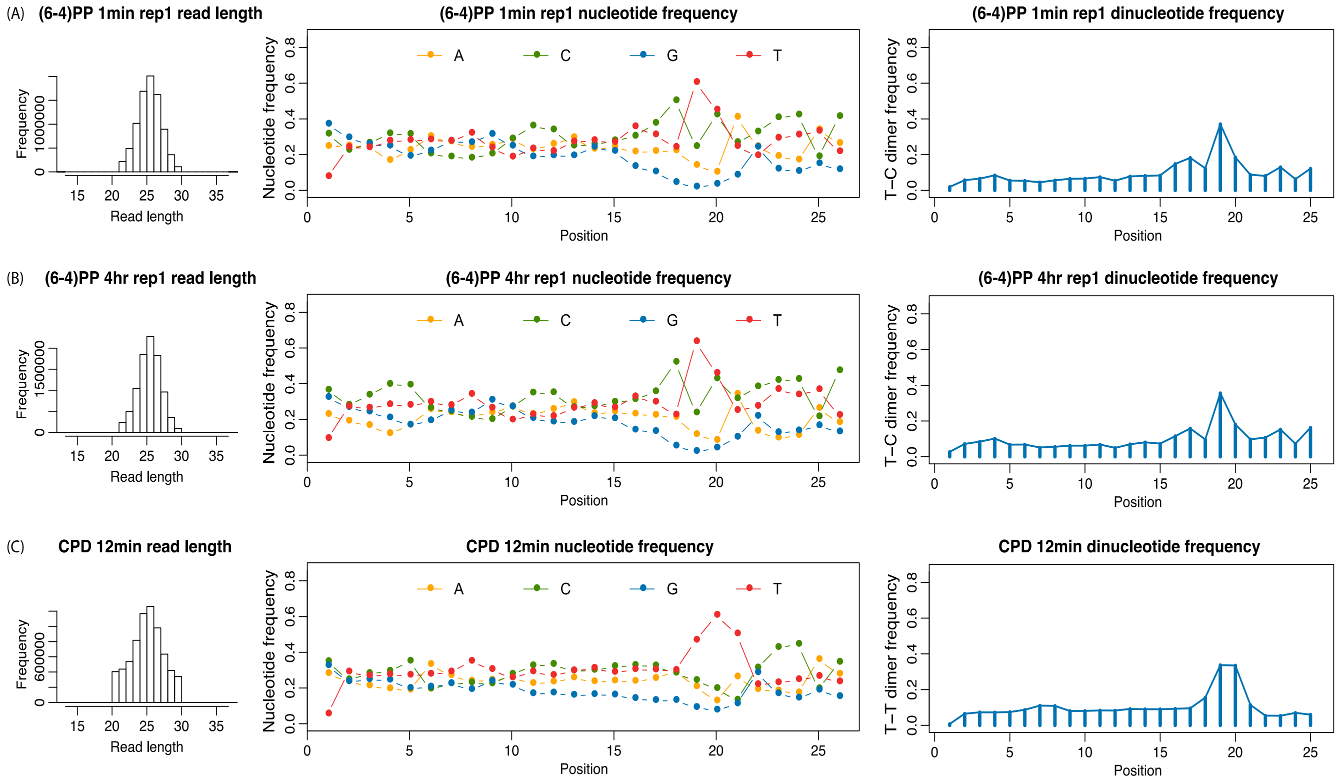

**Figure S3: Excision repair in transcribed strand (TS) versus non-transcribed strand (NTS).** For each gene, we quantify its strand-specific repair levels by XR-seq. Results are shown for (A) (6-4)PP XR-seq 1 min replicate 1, (B) (6-4)PP XR-seq 4hr replicate 1, and (C) CPD XR-seq 12 min. On the genome-wide scale, excision repair of damage on the TS and on the NTS is essentially equal, indicating no significant transcription-coupled repair of (6-4)PP at any timepoints and of CPD at the early 12 min timepoint.

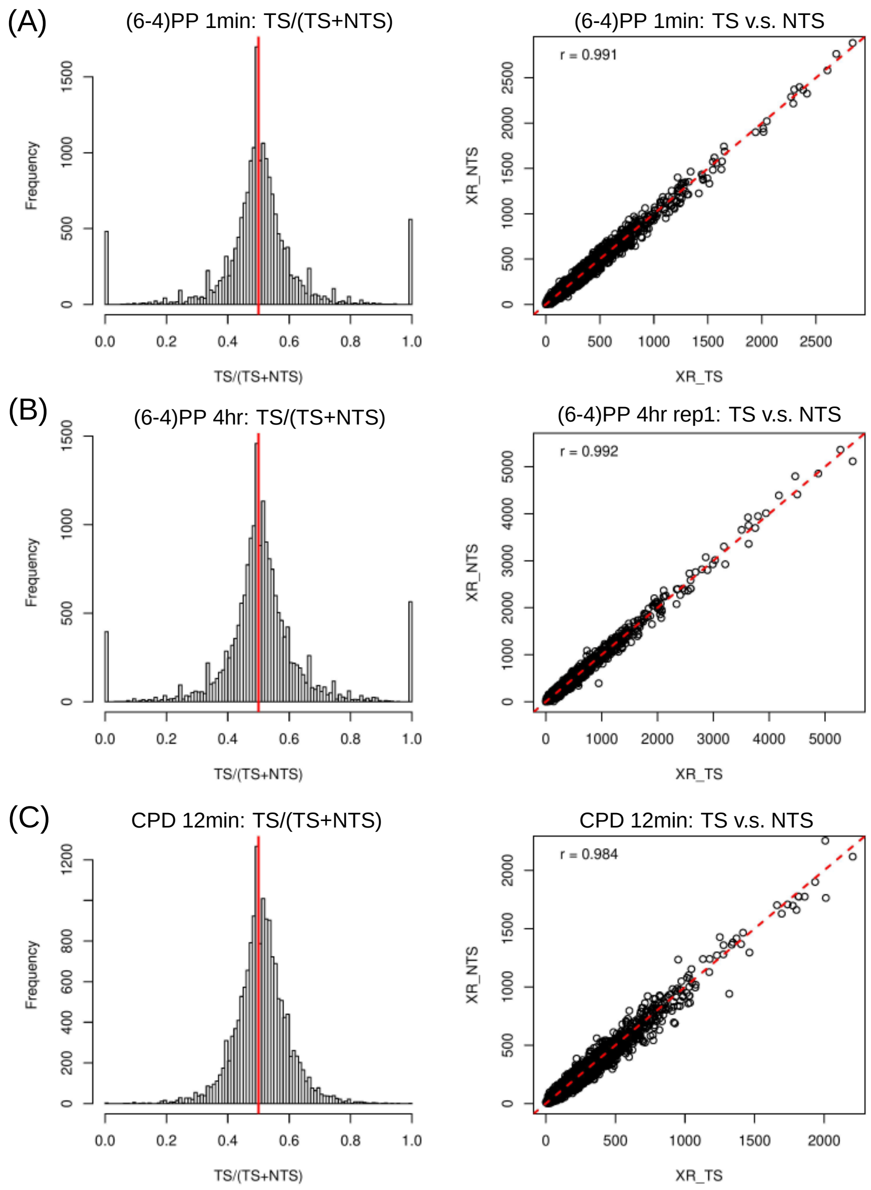

**Figure S5: Normalized *z*-scores from Poisson log linear model for CPD repair.** A cross-sample Poisson log linear model is applied to the repair data to account for library size difference, bin-specific artifacts, and TC content bias. The model is applied to the (A) plus and (B) minus strand separately for normalization. The detected repair hotspots and coldspots harbor a much higher level of the observed repair ($Y$) than that of the null repair as expected ($\lambda$), in the 1 min and 4 hr replicates, respectively.

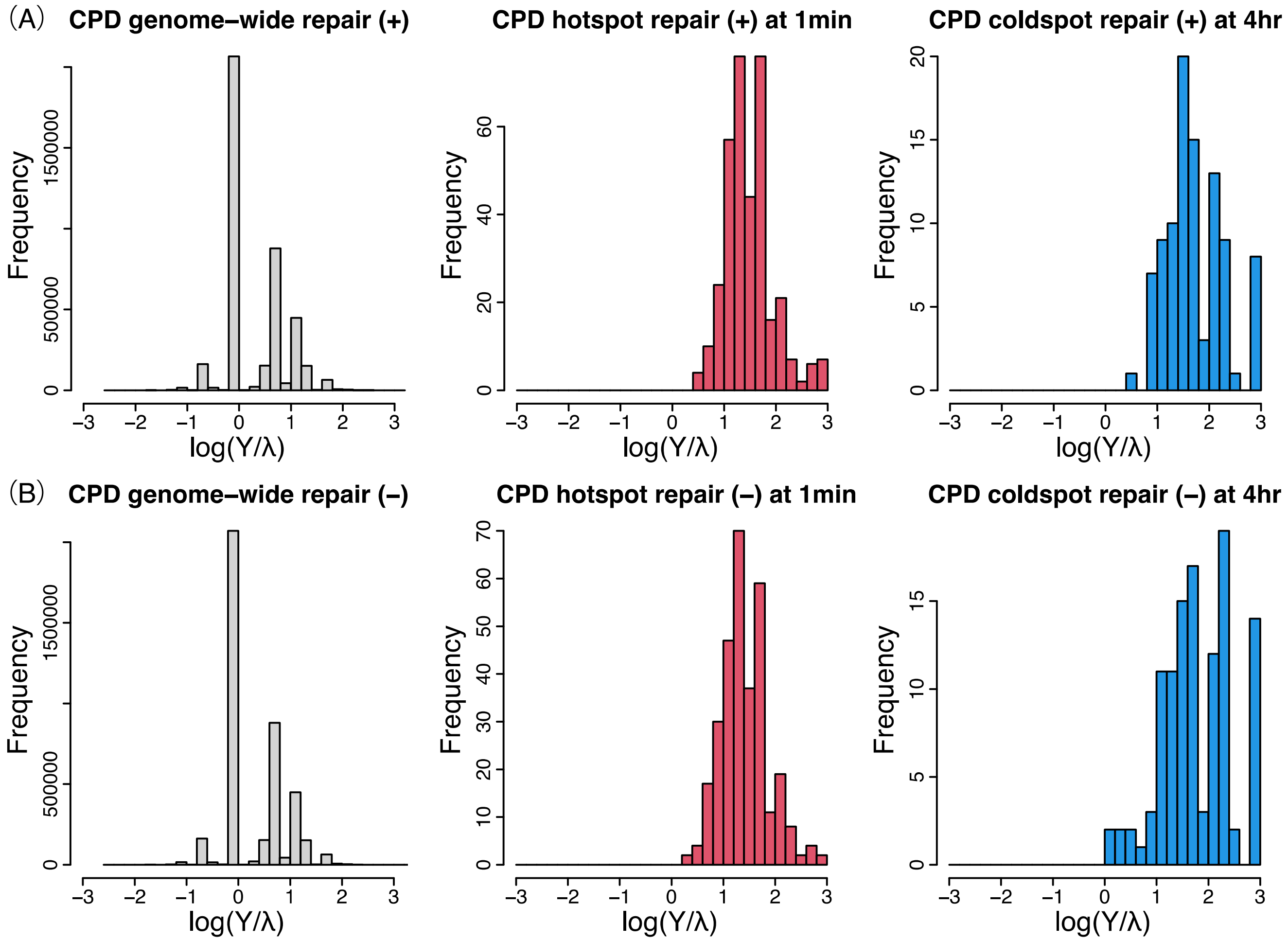

**Figure S6: Genome-wide distribution of identified repair hotspots and coldspots.** Repair hotspots for (6-4)PP and CPD, and repair coldspots for (6-4)PP are separated by strands and visualized on chromosomal ideograms using Phenogram[1]. The hotspots and coldspots are distributed across the entire genome fairly homogeneously.

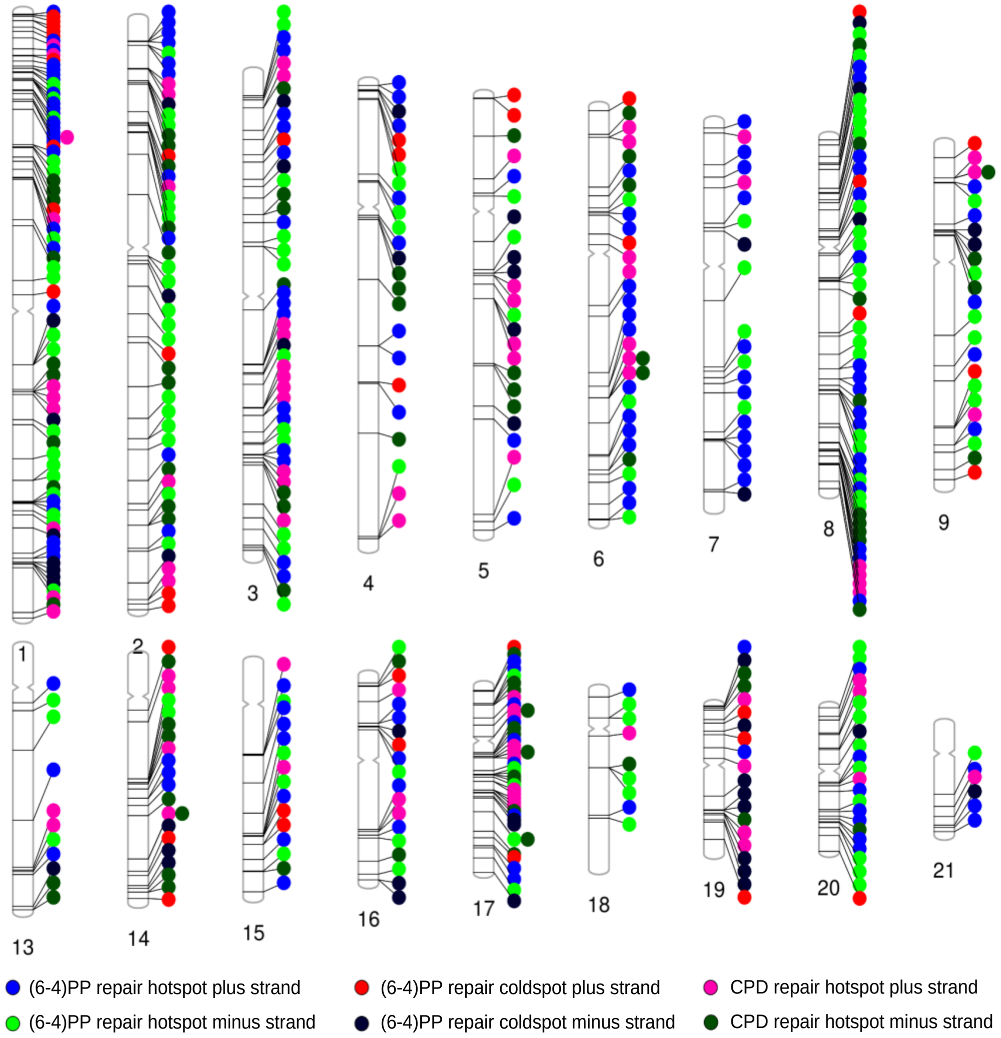

**Figure S7: Flanking regions of the identified hotspots and coldspots are enriched for cytosine.** Distributions of the nucleotide frequencies centered at the damage sites (marked as 0, 0) are shown. Frequencies are computed using all reads that are mapped to (A) (6-4)PP repair hotspots, (B) (6-4)PP repair coldspots, and (C) CPD repair hotspots. As a comparison, the same number of excision repair reads are randomly sampled from the genome and are used to generate the frequency distribution from the "random spot" in each panel.

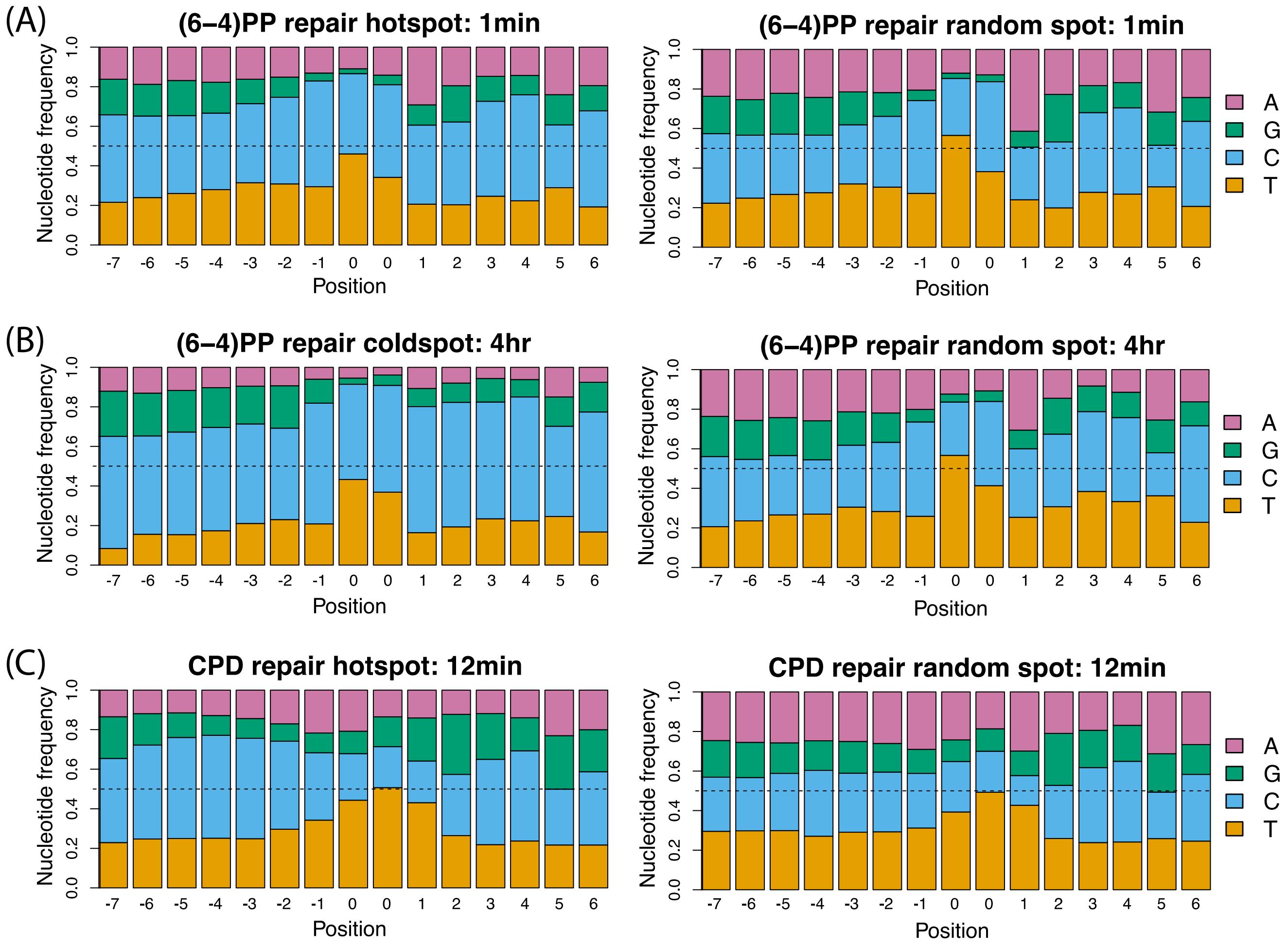

**Figure S8: Nucleotide frequency of cytosine in repair hotspots and coldspots compared to genomic average.** Histograms of percentage of cytosine are shown (A) across all genomic bins, (B) in (6-4)PP repair hotspots, (C) in (6-4)PP repair coldspots, and (D) in CPD repair hotspots. The identified repair hotspots and coldspots are enriched for cytosines in the flanking regions of the repair sites. Such observation is not due to sequence context biases—the identified repair hotspots and coldspots only take up a small fraction of the entire genomic regions that are cytosine-enriched.

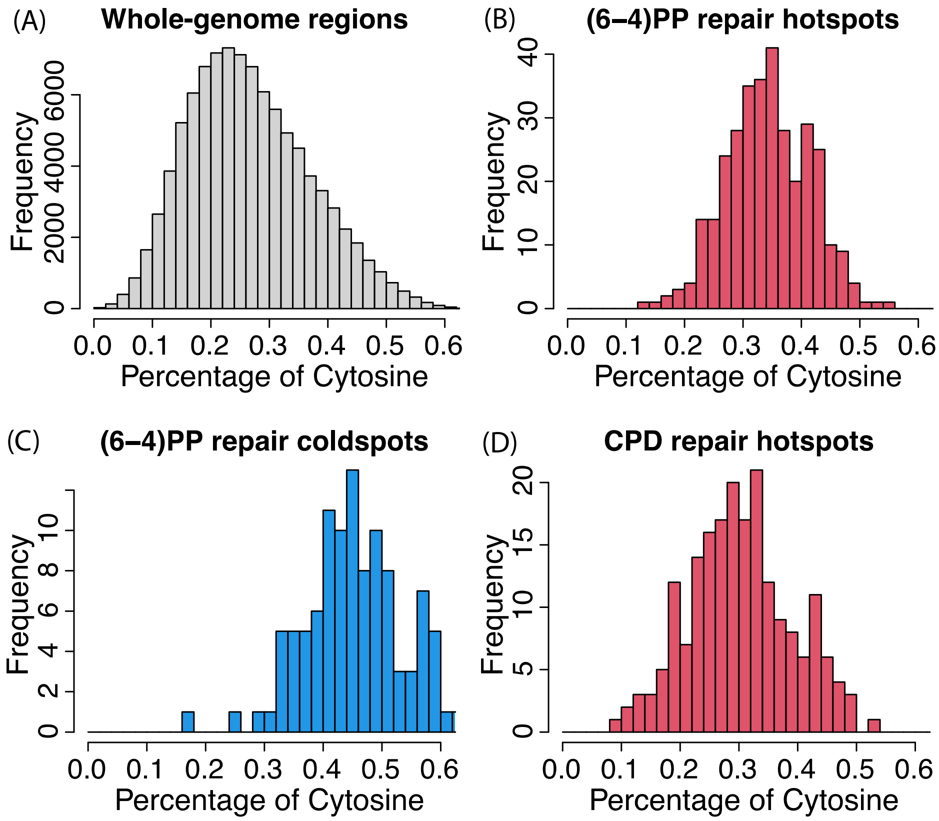

**Figure S9: Epigenomic markers and genome-wide annotations for CPD damage hotspots.** (A) DNase I hypersensitivity by DNase-seq and histone modifications by ChIP-seq bear no statistical differences between the CPD damage hotspots and random spots. (B) CPD damage hotspots are enriched for heterochromatin regions, a finding that is concordant with previous reports[2,3]. The damage hotspots are also enriched for inter-CpG and intronic regions, both of which are AT rich.

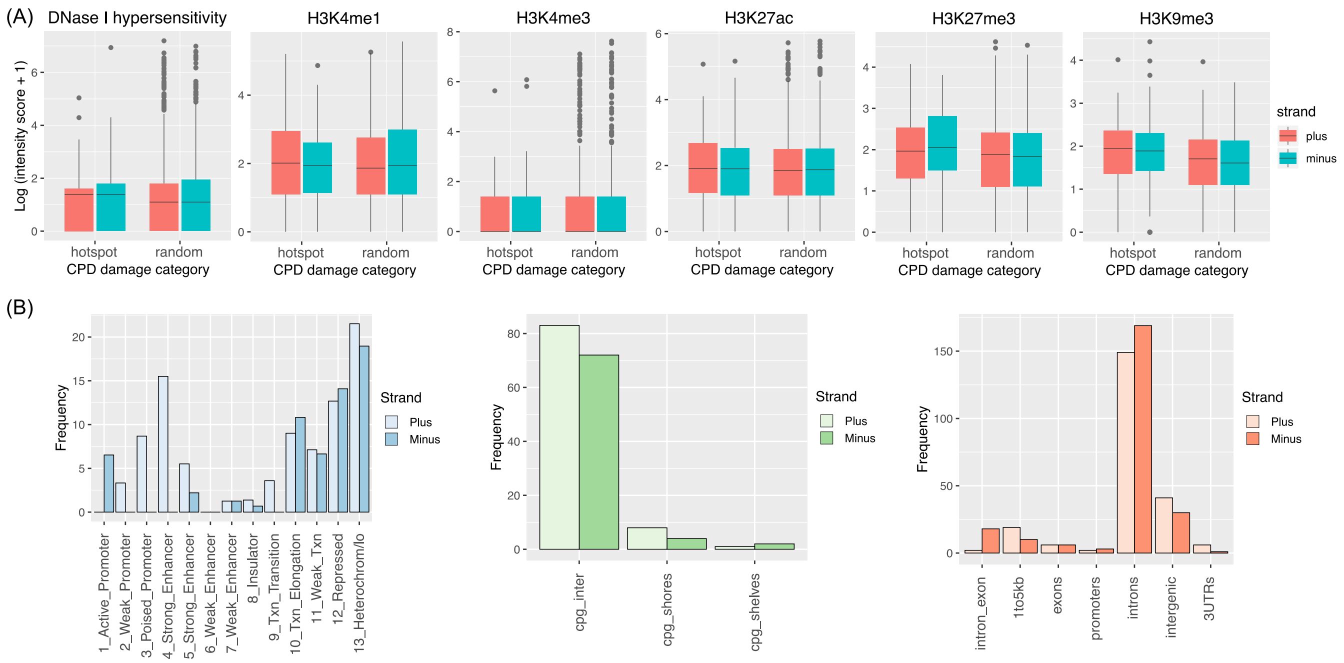

**Figure S10: Genome-wide repair kinetics across timepoints for strong enhancers, weak enhancers, repressed regions, and heterochromatin regions.** Enhancer regions are repaired at earlier timepoints, while repressed/heterochromatin regions are repaired at later timepoints. For strong enhancer and weak enhancer, the peaks at 5 min may be due to accumulation of non-degraded excision products from 1 min and 2 min.

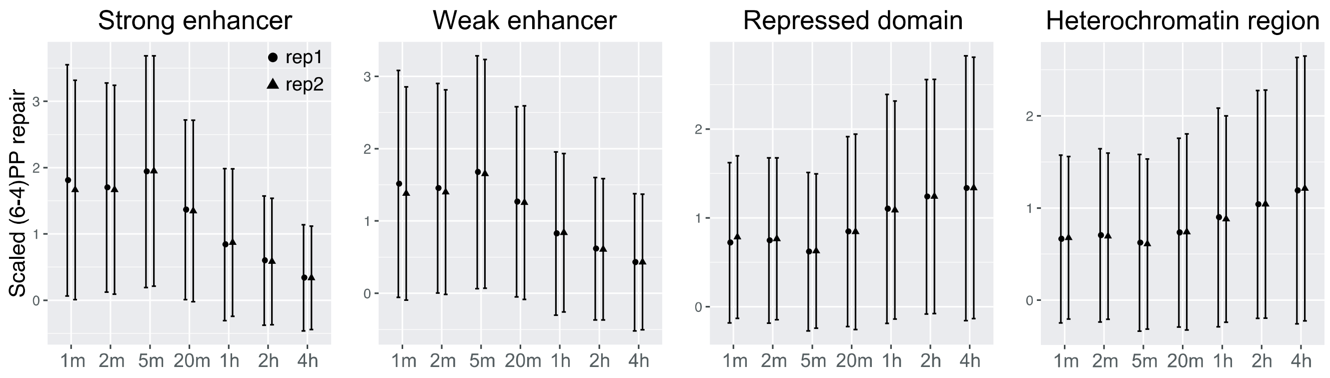

**Figure S11: Repair hotspots overlap with multiple consecutive enhancers from the same genomic regions.** (A) An example minus strand repair hotspot from chr 2. (B) An example plus strand repair hotspot from chr 1. Both hotspots overlap with super-enhancers.

(A)

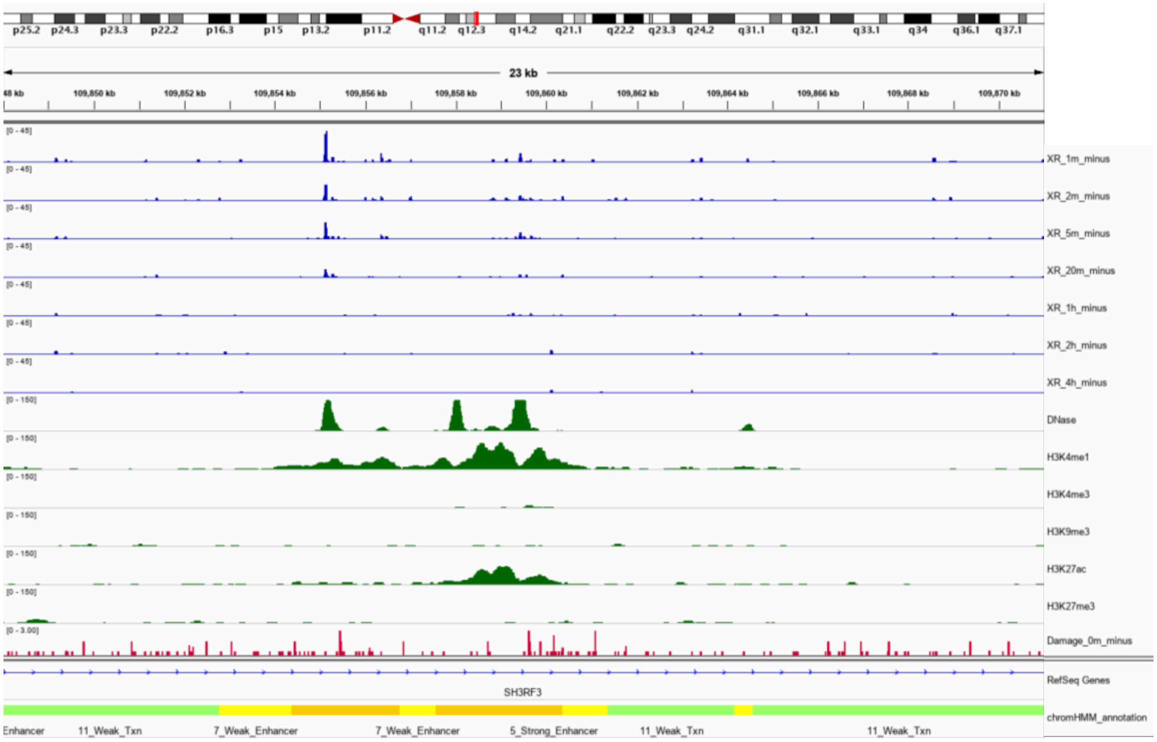


(B)

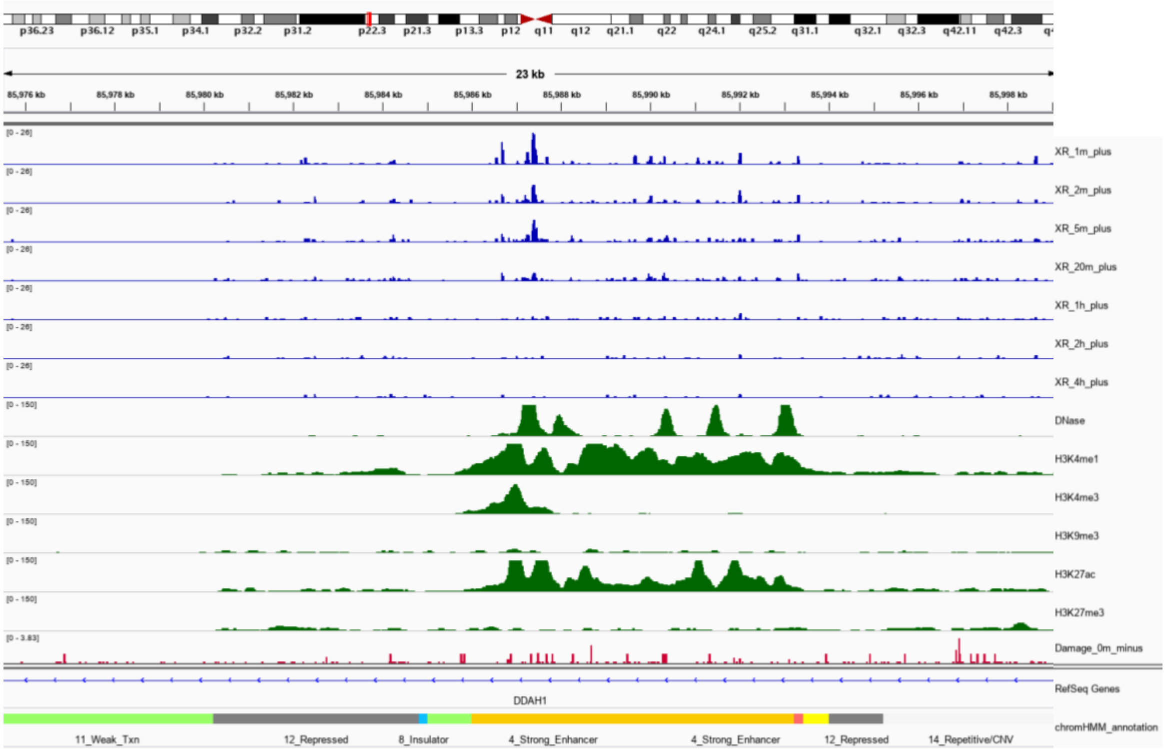

**Figure S12: Gene-specific repair kinetics.** (6-4)PP repair RPKMs as measured by XR-seq across different timepoints for genes from the circadian rhythm and excision repair pathways. TS and NTS repair are approximately on the same levels indicating no transcription-coupled repair at play. Different genes show different repair kinetics.

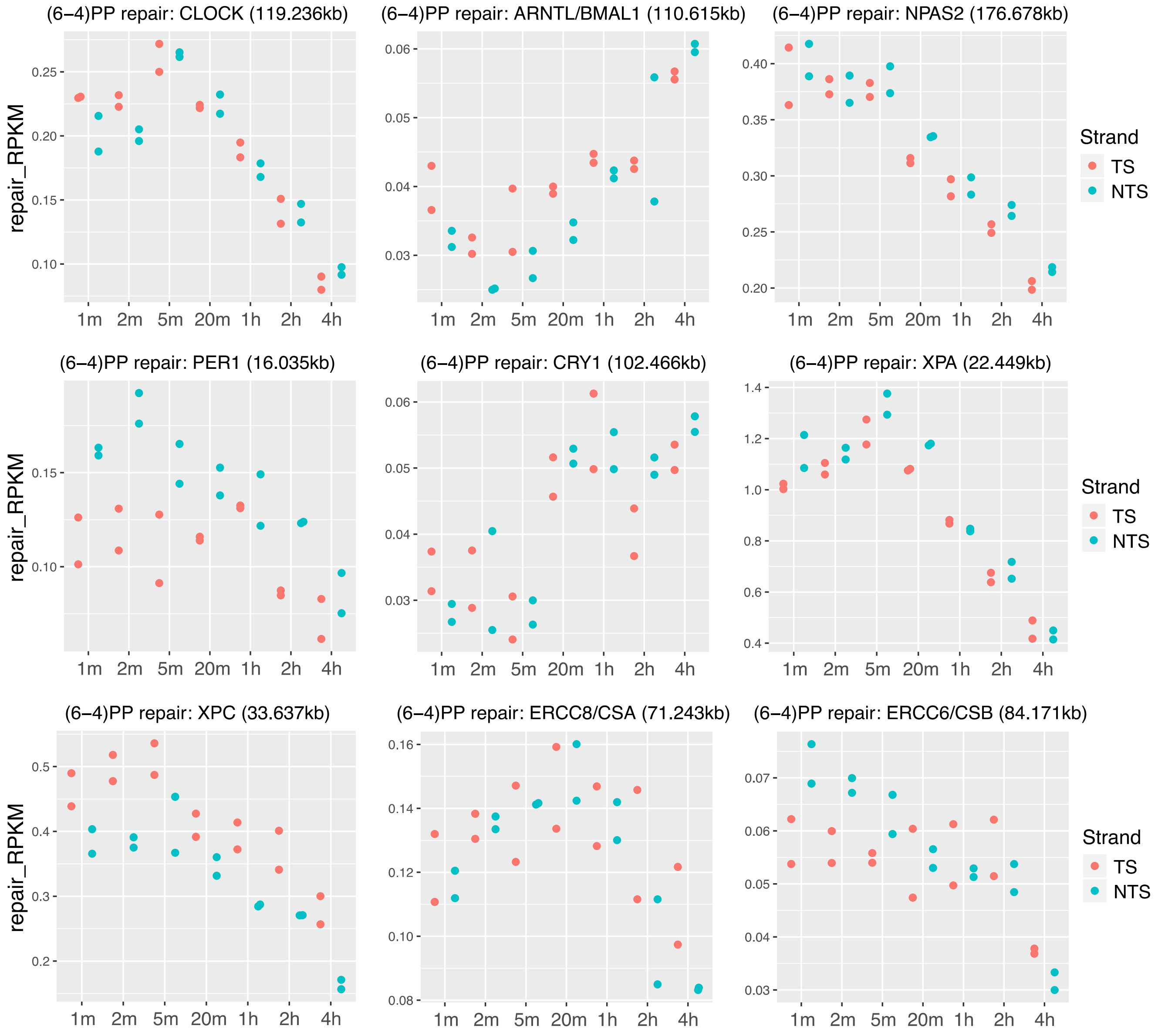

**Table S1: XR-seq sample information**. Number of reads is reported across all samples after each filtering step: fastq_reads is for the number of sequence reads in the raw sequencing file; total_mapped is for the total number of mapped reads; dedup is for the number of reads after deduplication; mapq is for the reads with mapping quality greater than 20; chr is for the reads that are mapped to the autosomes and the sex chromosomes; qwidth is the number of reads with lengths between 21 and 31bp; and genebody is the number of reads that are mapped to the gene bodies.

| fastq_name | antibody | treatment | time | replicate | fastq_reads | total_mapped | dedup | mapq | chr | qwidth | genebody |
|---|---|---|---|---|---|---|---|---|---|---|---|
| **N641MR1_CAGATC_S49_L008_R1_001.fastq** | 64 | 20J | 1m | 1 | 384802940 | 71402356 | 8401818 | 7941024 | 7930102 | 7930102 | 4326493 |
| **N641MR2_GATCAG_S50_L008_R1_001.fastq** | 64 | 20J | 1m | 2 | 422824212 | 67345807 | 8620110 | 8112862 | 8075112 | 8075112 | 4444832 |
| **N642MR1_ACTTGA_S51_L008_R1_001.fastq** | 64 | 20J | 2m | 1 | 279335280 | 53197655 | 23345765 | 22078323 | 22059768 | 22059768 | 11784233 |
| **N642MR2_TAGCTT_S52_L008_R1_001.fastq** | 64 | 20J | 2m | 2 | 248034148 | 45543788 | 16644732 | 15692341 | 15672060 | 15672060 | 8526503 |
| **N645MR1_ATCACG_L005_R1_001.fastq** | 64 | 20J | 5m | 1 | 104591936 | 19364359 | 16600527 | 15712745 | 15698665 | 15698665 | 8578429 |
| **N645MR2_TGACCA_L005_R1_001.fastq** | 64 | 20J | 5m | 2 | 82008780 | 15249676 | 12982483 | 12273842 | 12263361 | 12263361 | 6760420 |
| **N6420MR1_ACAGTG_L003_R1_001.fastq** | 64 | 20J | 20m | 1 | 146127236 | 26903762 | 20799106 | 19638269 | 19620576 | 19620576 | 10428629 |
| **N6420MR2_GATCAG_L003_R1_001.fastq** | 64 | 20J | 20m | 2 | 214152044 | 39984681 | 32266738 | 30521020 | 30494421 | 30494421 | 16161219 |
| **N641HR1_TAGCTT_L003_R1_001.fastq** | 64 | 20J | 1h | 1 | 194350240 | 34956862 | 30108510 | 28313194 | 28282892 | 28282892 | 14298448 |
| **N641HR2_GCCAAT_L003_R1_001.fastq** | 64 | 20J | 1h | 2 | 140819580 | 25008097 | 19975225 | 18762428 | 18741270 | 18741270 | 9587793 |
| **N642HR1_TAGCTT_L001_R1_001.fastq** | 64 | 20J | 2h | 1 | 60324552 | 10548055 | 9562799 | 8959193 | 8945613 | 8945613 | 4334201 |
| **N642HR2_TTAGGC_L001_R1_001.fastq** | 64 | 20J | 2h | 2 | 106209304 | 18167300 | 15697254 | 14698388 | 14679905 | 14679905 | 7102953 |
| **N644HR1_GGCTAC_L001_R1_001.fastq** | 64 | 20J | 4h | 1 | 60961524 | 10377267 | 9112994 | 8518484 | 8501855 | 8501855 | 3849252 |
| **N644HR2_TGACCA_L001_R1_001.fastq** | 64 | 20J | 4h | 2 | 99967420 | 16902160 | 13548187 | 12648710 | 12626997 | 12626997 | 5687242 |
| **NHF1CPD12minrep1.fastq** | CPD | 20J | 12m | 1 | 226271720 | 22029959 | 9193596 | 8146640 | 8146640 | 7002646 | 3659658 |

**Table S2: (6-4)PP repair hotspots.** Genomics coordinates and read depths for (A) 175 repair hotspots from the plus strand and (B) 156 repair hotspots from the minus strand. Separately attached as an excel file.

**Table S3: (6-4)PP repair coldspots.** Genomics coordinates and read depths for (A) 48 repair coldspots from the plus strand and (B) 57 repair coldspots from the minus strand. Separately attached as an excel file.

**Table S4: CPD repair hotspots.** Genomics coordinates and read depths for (A) 99 repair hotspots from the plus strand and (B) 93 repair hotspots from the minus strand. Separately attached as an excel file.

**Table S5: Motif analysis of identified repair hotspots and coldspots.** Motif analysis using the MEME suite[5] suggest enriched motifs with TC and TT dinucleotides at the damage/repair sites. For (6-4)PP repair hotspots and coldspots, a canonical sequence of CTCA is enriched, as expected. For all three categories, there is enriched cytosine in the flanking regions. The gene ontology hierarchy is specified by the abbreviations (BP for biological process, CC for cellular component, and MF for molecular function) and the name of the term.

| Category | Replicate | Sequence Count | Motif | Motif Sites | Top 3 Predictions |
|---|---|---|---|---|---|
| 6-4 repair hotspots | 1 | 6984 | 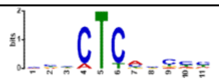 | 6652 | BP positive regulation of transcription from RNA polymerase II promoter<br>BP anterior/posterior pattern formation<br>MF calcium ion binding |
| | 2 | 5863 | 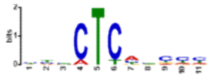 | 5561 | |
| 6-4 repair coldspots | 1 | 1004 | 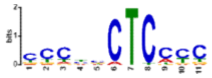 | 913 | CC transcription factor complex<br>MF potassium ion binding<br>MF transcription activator activity |
| | 2 | 990 | 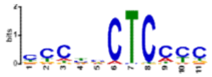 | 908 | |
| CPD repair hotspots | 1 | 2268 | 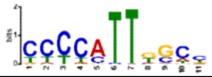 | 1241 | MF transcription factor activity<br>MF sequence-specific DNA binding<br>CC extracellular space |

**Table S6: CPD damage hotspots.** Genomics coordinates and read depths for (A) 91 damage hotspots from the plus strand and (B) 78 damage hotspots from the minus strand. Separately attached as an excel file.

**Table S7: Profiled FIRES, super-enhancers, and interactions using Hi-C data of human fibroblasts.** (A) FIREs calling results by FIREcaller[6] using Hi-C data of IMR90[7]. Outputs include continuous FIRE scores, dichotomous FIREs, and super-FIREs. (B) List of super-enhancers of human fibroblasts, inferred by Hi-C data of IMR90[8]. (C) Significant interactions inferred from the Hi-C data[9]. Separately attached as an excel file.

**Table S8: Repair hotspots and coldspots overlapped with FIREs, super-enhancers, and significant interactions.** Number of overlapped FIREs, super-enhancers, and significant interactions for (A) (6-4)PP repair hotspots, (B) (6-4)PP repair coldspots, and (C) CPD repair hotspots. Separately attached as an excel file.

**Table S9: Genomic segments for replication timing of human fibroblasts.** Results are aggregated from a re-analysis[10] of the Repli-Seq data of human fibroblasts[11]. Separately attached as an excel file.

**Table S10: Segmented regression analysis results of the time-course (6-4)PP repair data.** Results were generated by Trendy[4]. Separately attached as an excel file.

**Table S11: Significantly enriched gene ontology (GO) biological processes for significant genes returned by Trendy.** Genes with adjusted $R^2$ greater than 0.8 are included in the GO analysis using PANTHER[12]. Analysis was carried out separately on four groups of genes with (A) down, (B) up-down, (C) up, and (D) down-up fitted trend. Separately attached as an excel file.
